# Neutral transcriptome rewiring promotes QDR evolvability at the species level

**DOI:** 10.1101/2024.02.15.580486

**Authors:** Florent Delplace, Mehdi Khafif, Remco Stam, Adelin Barbacci, Sylvain Raffaele

## Abstract

Quantitative disease resistance (QDR) is an immune response limiting pathogen damage in plants. It involves transcriptomic reprogramming of numerous genes, each having a small contribution to plant immunity. Despite QDR broad-spectrum nature, the evolution of its underlying transcriptome reprogramming remains largely uncharacterized. Here, we analyzed global gene expression in response to the necrotrophic fungus *Sclerotinia sclerotiorum* in 23 *Arabidopsis thaliana* accessions of diverse origin and contrasted QDR phenotype. Over half of the species pan-transcriptome displayed local responses to *S. sclerotiorum*, with global reprogramming patterns incongruent with accessions phylogeny. Due to frequent small-amplitude variations, only ∼11% of responsive genes were common across all accessions, defining a core transcriptome enriched in highly-responsive genes. Co-expression and correlation analyses showed that QDR phenotypes result from the integration of numerous genes expression. Promoter sequence comparisons revealed that variation in DNA-binding sites within cis-regulatory regions contributing to gene expression rewiring. Finally, transcriptome-phenotype maps revealed abundant neutral networks connecting diverse QDR transcriptomes with no loss of resistance, hallmarks of robust and evolvable traits. This navigability associated with regulatory variation in core genes highlights their role in QDR evolvability. This work provides insights into the evolution of complex immune responses, informing models for plant disease dynamics.

**Classification**: Biological Sciences, Plant Biology

## Introduction

Plants actively respond to pathogen attacks by modulating their physiology, using a range of molecular mechanisms to detect pathogens and defend against them. Gene-for-gene resistance is an immunity mechanism which involves single dominant resistance genes belonging to the nucleotide-binding domain and leucine-rich repeat (NLR) or Receptor Like Protein (RLP) families (Dodds and Rathjen, 2010). Quantitative disease resistance (QDR) is another form of plant immunity that limits the damage caused by pathogen infection which is nearly universal, often durable and broad-spectrum (Roux *et al*., 2014; Corwin and Kliebenstein, 2017). QDR involves genes from very diverse families, each having a small effect on resistance (Gou *et al*., 2023), which makes the identification and characterization of QDR genes difficult. Interestingly, QDR is the only form of plant immunity that reduces disease symptoms caused by many organisms that actively kill host cells for infection, such as the necrotrophic pathogen *Sclerotinia sclerotiorum* (Niks *et al*., 2015; Wang *et al*., 2019). This devastating pathogen is responsible for white and stem mold diseases on a broad range of dicot plants (Bolton *et al*., 2006), and notably causes severe damage on Brassicaceae, including *Arabidopsis thaliana*, when conditions are favorable (Derbyshire and Denton-Giles, 2016).

QDR involves reprogramming of a large number of genes (Pink *et al*., 2023; Zhang *et al*., 2019; Wu *et al*., 2016; Sucher *et al*., 2020). Their individual contribution to the resistance phenotype is largely unknown but is likely low, in agreement with an infinitesimal model of complex polygenic traits (Nelson *et al*., 2013; Boyle *et al*., 2017). Yet, stress-responsive genes and the transcription factors regulating them may play crucial roles in plants adaptation to pathogen attacks (Coolen *et al*., 2016; Shankar *et al*., 2016; Groen *et al*., 2020). Understanding how plants adapt their transcriptomes to mitigate damage caused by necrotrophic pathogens is crucial to understand the evolutionary dynamics of complex traits at the transcriptome level, and to document the diversity and functioning of QDR genetic determinants.

Upon pathogen attack, plant lineages display diversified metabolic and transcriptomic responses that converge on essential hubs (Han and Tsuda, 2022). Convergent pathways, exemplified by the intricate hormone signaling network, drive the expression of core genes across long evolutionary timescales (Aerts *et al*., 2021) and buffer metabolic and transcriptomic plasticity of plant defense (Zhang *et al*., 2017). The evolution of the plant immune system is determined by variation both in coding sequences and in gene expression (Tsuda and Somssich, 2015; Monteiro and Nishimura, 2018). Local diversity in gene expression pattern across species and populations can be responsible for adaptive evolution (Harrison *et al*., 2012; Lasky *et al*., 2014). Several studies compared resistant with susceptible plant genotypes (Westrick *et al*., 2019; Mwape *et al*., 2021) or used expression Quantitative Trait Loci (eQTL) (Moscou *et al*., 2011) to point at genomic loci that control gene-expression differences and identify determinants and regulators of *S. sclerotiorum* resistance. However, studies on global expression diversity in response to pathogens at the species level remain scarce, and the extent to which diverse patterns of transcriptome reprogramming can lead to similar QDR phenotype is unclear. Transcriptional differences within species primarily rely on DNA sequence variations, including single nucleotide polymorphisms (SNPs), insertions or deletions (indels), and transposable elements (TEs) (Studer *et al*., 2011; Ando *et al*., 2018; Soltis *et al*., 2020). Variation in transcription factor (TFs) binding motifs in 5′-regulatory regions modulate transcriptomic defense responses, such as in Brassicaceae where the emergence of WRKY TF binding motifs associates with specific response to the bacterial molecular pattern flg22 (Winkelmüller *et al*., 2021). Cis-regulatory variantscould be a major source of transcriptome adaptation during plant evolution by acting on TFs binding sites (Filichkin *et al*., 2011; Yu *et al*., 2015; Wang *et al*., 2016), but their contribution to QDR variation at the species level is currently elusive.

Quantitative resistance is often linked to long-term durability, and partial resistance tends to be prevalent in plant populations (Corwin and Kliebenstein, 2017). However, the mechanisms driving the emergence, maintenance, and evolutionary dynamics of QDR transcriptome reprogramming remain poorly understood. According to neutral evolution theories, stochastic processes may play a significant role in shaping transcriptome evolution, particularly in plants (Broadley *et al*., 2008). One of the key challenges is predicting evolutionary outcomes within this neutral framework, which posits that diverse genotypes can converge to produce similar phenotypes (Nimwegen et al., 1999). Conserved transcriptome profiles suggest that selective pressures, such as purifying or positive selection, help to maintain beneficial transcriptomic patterns while avoiding deleterious phenotypes (Rifkin *et al*., 2005; Melo and Marroig, 2015; Melo *et al*., 2016). The plasticity of transcriptome reprogramming, facilitated by mutations in both cis- and trans-regulatory elements, may be a key driver of expression variation within and between species (Zheng et al., 2011; Hill et al., 2021). Understanding the accessible evolutionary pathways and how they are navigated in the context of QDR could provide valuable insights on both natural evolutionary dynamics and potential avenues for directed evolution.

Despite the widespread nature of QDR and its importance for the durable management of plant diseases, our understanding of QDR evolution is limited to knowledge on specific genes or gene families that were associated to this form of immunity (Badet *et al*., 2017; Barco *et al*., 2019). The conservation of global QDR responses at the species level offers a unique opportunity to study the recent evolution of gene expression patterns underlying complex traits such as QDR. In this paper, we explored the intraspecific variation in *A. thaliana* transcriptomic responses to *S. sclerotiorum* across 23 accessions with diverse resistance phenotypes and geographical origins. We identified a set of ∼2000 genes strongly responsive to *S. sclerotiorum* inoculation and under purifying selection, thus defining the *A. thaliana* core transcriptome. Cross-accession comparison of promoter sequences highlighted recent presence/absence polymorphisms in cis-regulatory elements associated with variations in modules of co-expressed genes and genes whose expression correlated with QDR. Finally, we constructed transcriptome-phenotype maps relating global transcriptome reprogramming and QDR at the species level to explore the evolution of this trait. We reveal abundant neutral networks connecting diverse transcriptomes with no loss of disease resistance. These landscapes support a role for the core transcriptome in the robustness and evolvability of QDR. These findings advance our understanding of the evolutionary dynamics of complex immune responses at the species level.

## Materials and Methods

### Plant material and RNA sampling

*Arabidopsis thaliana* accessions listed in Table S1 were obtained from The Versailles Arabidopsis Stock Center. General information regarding collection location and climate data of these accessions was retrieved from https://1001genomes.org and https://www.worldclim.org. ADMIXTURE genetic groups corresponding to the 23 accessions were recovered from (Alonso-Blanco *et al*., 2016). Plants were grown in Jiffy pots under controlled conditions at 22°C with a 9-hour light period at an intensity of 120 μmol/m^2^/s. Five-week-old plants were inoculated with *S. sclerotiorum* strain 1980 (ATCC18683) on three leaves using 0.5-cm-wide plugs of Potato Dextrose Agar (PDA) medium colonized by the fungus. The agar plugs containing the fungal pathogen were placed on the adaxial surface of leaves, and plants were incubated in trays sealed with plastic wrap to maintain 80% relative humidity. 24 hours after inoculation, leaf rings including the periphery of disease lesions were collected as in (Peyraud *et al*., 2019). Material obtained from three leaves per plant were pooled as one sample, samples were collected in three to five replicates. Leaves from non-inoculated plants under the same conditions were collected as control at the same time.

### RNA extraction, sequencing and mapping

Samples were ground using glass beads (2.5 mm) and RNA was extracted using NucleoSpin RNA PLUS extraction kits (Macherey-Nagel) following the manufacturer’s protocols. The quality of the extracted RNA was assessed using Agilent bioanalyzer nanochips. Libraries preparation and mRNA sequencing was conducted at the GeT-Plage facility (INRAE Castanet Tolosan, France) using the Illumina TrueSeq Stranded mRNA kit, 2×150 base pair reads sequencing with S4 chemistry on a NovaSeq instrument. For each experimental condition (inoculated or non-inoculated), three to five independent biological samples were processed, and their respective Illumina paired-end reads were obtained (Table S2). These reads were simultaneously aligned to the Arabidopsis Columbia-0 reference genome Araport11, and the *S. sclerotiorum* strain 1980 version 2 genome (Derbyshire *et al*., 2017), using the pipeline nf-core/rnaseq version 3.12.0 described in https://doi.org/10.5281/zenodo.1400710.

### Identification and analysis of Differentially Expressed Genes

Normalization of read counts per gene was performed using the edgeR package considering total read count by library (Robinson *et al*., 2010). Differential expression analysis was conducted using edgeR package v 3.32.1 in R 4.0.4. using mock treated plants as a reference for each accession with ∼replicates + treatment as the design formula. Genes with |Log2 Fold Change (LFC)|≥ 2 and a Benjamini–Hochberg adjusted P-value < 0.05 were considered significant for differential expression. Raw and processed gene expression data was deposited in NCBI’s GEO under accession number GSE248079. Heatmaps were generated using the heatmap.2 package in R 4.2.1. Hierarchical clustering was performed using the default parameters of hclust, and the robustness of the branches was evaluated with pvclust v2.2-0. Gene Ontology enrichment analysis was performed using BINGO module 3.0.5 from Cytoscape 3.9.1, assessing over-representation of GO terms in our gene lists compared all GO terms from *A. thaliana*.

Log regression analyzes to extrapolate the number of responsive and core genes were done using ’drc’ package v3.0-1 in R, the log-logistic (LL.4) model was employed to characterize the relationship between the number of DEGs (’n_genes’) and categorical grouping factors (’n_accessions’). The maximal gene expression level was estimated at 10^12^ for ’n_accessions.’. Visualizations were created using the ’ggplot2’ package, showcasing boxplots and overlaid red curves representing the fitted exponential trends.

### Phenotypic Characterization of Arabidopsis Accessions in response to *S. sclerotiorum*

0.5-cm-wide plugs of PDA agar medium containing *S. sclerotiorum* strain 1980 grown for 72 hours at 20°C on 14 cm Petri dishes were placed on the adaxial surface of detached leaves. These leaves were positioned in Navautron system (Barbacci *et al*., 2020), records were done using high-definition (HD) cameras “3MP M12 HD 2.8-12mm 1/2.5 IR 1:1.4 CCTV Lens”. In total 33 to 135 leaves from 3 experiments were inoculated per accession. Kinetics of *S. sclerotiorum* disease lesions were analyzed using INFEST script v1.0 (https://github.com/A02l01/INFEST). Statistical analyses of disease phenotypes were conducted using Tukey test in R 4.2.1 as described in (Barbacci *et al*., 2020). Lesion doubling time in minutes was calculated using the formula 10 * ln(2) / β, where β is the speed obtained with the INFEST script v1.0.

### Phylogenetics tree construction

Phylogenetic trees of the accessions were constructed from all coding sequences (CDS) recovered from the 23 corresponding pseudo-genomes available at the 1001 Genome Project Data Center. For this, pseudo-genomes were annotated using *A. thaliana* functional annotations (TAIR version 10) with liftoff (v1.6.1) (Shumate and Salzberg, 2021). Annotations were converted to coding sequences and protein sequences with gffread (Pertea and Pertea, 2020). 21,787 orthology groups were inferred with Orthofinder version 2.5.4 (Emms and Kelly, 2019). Orthogroups with a unique copy in each accession were aligned with Mafft (v7.313) (Nakamura *et al*., 2018). A super alignment was built from the orthogroup alignment by Orthofinder. Accession trees were inferred with IQ-TREE (v2.5.4) (Nguyen *et al*., 2015). The best models according to the Bayesian Information Criterion (BIC) computed by ModelFinder (Kalyaanamoorthy *et al*., 2017) were VT+F+I+G4 and GTR+F+I+G4 for the accession tree computed from protein alignments and from coding sequence alignments respectively. Branch support values were computed by two different methods: SH-aLRT (Guindon *et al*., 2010) and UFBoot (Hoang *et al*., 2018). Accession clustering was done using the cophenetic function from the R package ape v5.7.1. The tanglegram in Figure S2 was generated using phylogenetic tree data and the DEGs heatmap accession dendrogram using cophylo function from phytools v2.1-1.

### Ortholog Identification and Population Genetics Metrics in Arabidopsis

Arabidopsis orthologous gene groups were retrieved using the Phytozome v13 database (Goodstein *et al*., 2012). These orthogroups were used to assign relative gene age (from the Arabidopsis thaliana node to the Viridiplantae node), based on the species tree node to which the MRCA (Most Recent Common Ancestor) of the species represented in the orthogroup maps.. Enrichment of gene sets in gene age categories were analyzed with Chi-square tests at a significance threshold P-value < 0.05.

To investigate genetic diversity at the species level across 23 accessions, population genetics analyses were conducted using Variant Call Format (VCF) files obtained from https://1001genomes.org. These files were processed using Python version 3.8.19, with data handling and result exporting performed using Pandas (version 2.0.3). Scikit-allel (version 1.3.7) was employed to calculate population genetics metrics, including nucleotide diversity (π), Watterson’s Theta (θW), and Tajima’s D. Genetic variants among the VCF files were annotated using SnpEff v5.2c to classify variants as synonymous (no amino acid change) or non-synonymous (amino acid change) based on the Araport11 Arabidopsis annotation. To extract mutation types, the annotated VCF file was parsed with SnpSift, which facilitated the identification of synonymous and non-synonymous mutations based on variant effect annotations. BCFtools (version 1.2) was utilized to manage the VCF files and analyze trait variants, Vcftools (version 0.1.16) was employed to calculate nucleotide diversity for non-synonymous mutations (πN) and synonymous mutations (πS). Statistical analyses were performed using 1,000 bootstrap replicates of randomly sampled subsets with the same sample size. A Dunn’s test was conducted for each sampling, and a significant difference in at least 950 out of 1,000 samples was considered statistically significant.

### Regression analysis, disease score correlation with expression for each gene

To identify genes whose expression correlates with disease phenotype within accession subsets, the mean disease index by accession (the growth speed of the fungus refered as speed hereafter) was fitted with log fold change (LFC) of expression for each gene (hereafter ‘LFC’) in corresponding accessions by linear regression. We utilized all LFC data, taking into account the 21/23 LFC values with the lowest false discovery rate (FDR) values. We employed R2 and RMSE as evaluation metrics. R2 quantifies the proportion of variation in dependent variables that can be accounted for by the predictors, while RMSE represents the model’s average error. R2 and RMSE were calculated in R using ggplot2 v3.4.2 with the following formulas: R2 = (cor(speed, LFC))^2, and rmse = sum((speed - cov(speed, LFC) / var(LFC) * LogFC - mean(speed) - cov(speed, LFC) / var(LFC) * mean(LFC))^2), a rmse threshold of 0.05 was used. Based on R2 > 0.4 and RMSE < 0.05, thirty-three genes fitting these characteristics were identified. Due to unclear annotation of the promoter region of AT4G31405 in all accessions except Col-0, this gene was removed from our analysis.

### Promoter sequence analyses

500pb promoter sequences were retrieved from for the 23 accessions using gft files generated by liftoff (v1.6.1) (Shumate and Salzberg, 2021) for phylogenetic tree construction. To conduct DAP cis-motif enrichment analysis, we employed MEME Suite 5.5.7 (Bailey *et al*., 2015) with the Simple Enrichment Analysis (SEA) tool, using the Arabidopsis DAP (DNA Affinity Purification) motifs database from (O’Malley *et al*., 2016). For the identification of DAP cis-motifs predicting gene expression variation, we used the Boruta feature selection algorithm (Boruta v.8.0.0) implemented in R (v.4.0.5) with significance threshold P-value < 0.05. Tentative features were resolved using the TentativeRoughFix method. Gene-specific datasets were prepared by extracting LFC values and DAP cis-regulatory motif data from 500 bp promoter regions, identified using the Find Individual Motif Occurrences (FIMO) tool, based on the complete DAP motifs database from O’Malley *et al*. (2016). The Boruta algorithm was applied to assess the importance of motifs in explaining LFC variation across accessions. Final feature importance was determined by the mean importance score and the algorithm’s final decision for each motif.

### Comparative analysis of transcriptome similarity among accessions

Pearson correlation matrix to correlate transcriptomes across accessions, was performed using the cor function excluding auto-comparisons from stats package v4.3.1, considering the log fold change (LFC) for all expressed genes. The resulting matrix was subjected to hierarchical clustering using the pheatmap library v1.0.12 in R, with a specified number of clusters (k=4) and the “ward.D” method. The same clustering pattern was obtained using a distance matrix with dist function from stats package v4.3.1.

### Co-expression genes modules identification and analysis

We employed Weighted Gene Co-expression Network Analysis (WGCNA) (R package WGCNA, version 1.75-5) to identify gene co-expression modules across 23 accessions. To cluster genes with similar patterns of differential expression changes and to directly compare expression changes between accessions, we used the log fold change (LFC) data of all expressed genes. Hierarchical clustering was performed on the Topological Overlap Matrix (TOM), and dynamic tree cutting (R package dynamicTreeCut, version 1.63-1) was used to identify co-expression modules. Finally, the UMAP dimensionality reduction technique (R package umap, version 0.2.10.0) was applied to visualize the module structure. We calculated Spearman correlations between dendrogram-derived distance matrices (from gene expression clustering) and phenotypic distance matrices. For each module, gene expression data were clustered using hierarchical clustering (Euclidean distance, complete linkage), and the resulting cophenetic distance matrix was compared to phenotypic distance using a Spearman permutation test (n = 9,999)

### Transcriptome-phenotype map construction and navigability

Transcriptome-resistance maps were generated using the two major principal components of PCA analysis (prcomp function, stats package v4.3.1), based on all LFC values from the 23 accessions. The x-axis corresponds to PC1, and the y-axis corresponds to PC2. Resistance values (speed) for each accession were added as the third dimension on the z-axis. For representational purposes, resistance was calculated based on the fungus’s propagation rate. To maintain consistency in units, resistance was computed for each accession as the absolute difference between the progression rate and the maximum value of this rate measured for the given accession. These scattered data were then adjusted using multilevel B-Splines to reconstruct the transcriptome-resistance maps (R package MBA: Multilevel B-Spline Approximation version 0.1-0). To assess the navigability associated with an accession, we estimated the ratio of the transcriptome-resistance map area covered by all trajectories starting from the given accession. The navigable area was obtained using a constrained random walker, with the starting point corresponding to the accession’s position on the map. At each iteration, the constrained random walker could move randomly to any of the 8 adjacent positions, provided the trajectory leads to a position with at least equal resistance. When the constrained random walker could no longer move, the length of its trajectory was recorded. A new trajectory was then estimated by resetting the walker to its initial position. This process was repeated 5,000 times for each accession. The navigability associated with the accession was then calculated as the union of all trajectories, normalized by the total surface area of the map.

## Results

### QDR phenotype and expression similarity are decorrelated in *A. thaliana* admixture groups

To explore the evolution of *Sclerotinia sclerotiorum* resistance at the species level, we collected 23 natural *A. thaliana* accessions covering a broad range of genetic diversity, geographic distribution, and climatic origins (Figure 1A, Figure S1). A phylogenetic tree based on CDS derived from pseudogenomes identified 8 of the 9 major ADMIXTURE genetic ancestry groups, or genetic clusters described in Alonso-Blanco *et al*., 2016. To relate genetic origin with disease resistance variation at the species level, we evaluated the susceptibility to *S. sclerotiorum*. Susceptibility ranged from a mean lesion doubling time of 306.7 minutes (Se-0) to 103.9 (Co-1), representing approximately a 3-fold increase in susceptibility (Figure 1B). The clusters included accessions with contrasted susceptibility levels, indicating that QDR varied largely independently within the groups. To document *A. thaliana* species pan-transcriptome diversity, we performed RNA-sequencing in 23 accessions. To assess the global gene expression profile during infection, total mRNA of healthy leaves and leaves colonized by *S. sclerotiorum* were collected 24 hours post-inoculation (hpi) and sequenced. Each sample was collected in independent triplicates to allow the detection of accession-specific differentially expressed genes (DEGs). We obtained an average 32,908,875 reads per sample, 99% of which could be mapped to Araport11 reference transcripts in healthy leaves and 43% in infected leaves (Table S3). A consistent 60.92% ± 1.72% of complete transcriptomes was expressed across the 23 accessions in healthy plants, and 54.71% ± 3.51% in infected plants. To identify *S. sclerotiorum*-responsive transcripts, we determined DEGs in infected versus healthy leaves for each accession (Figure S2). The number of DEGs ranged from 6,712 to 9,269, with a majority of DEGs downregulated in every accession (upregulated/downregulated ratio of 0.72

**Figure 1:**
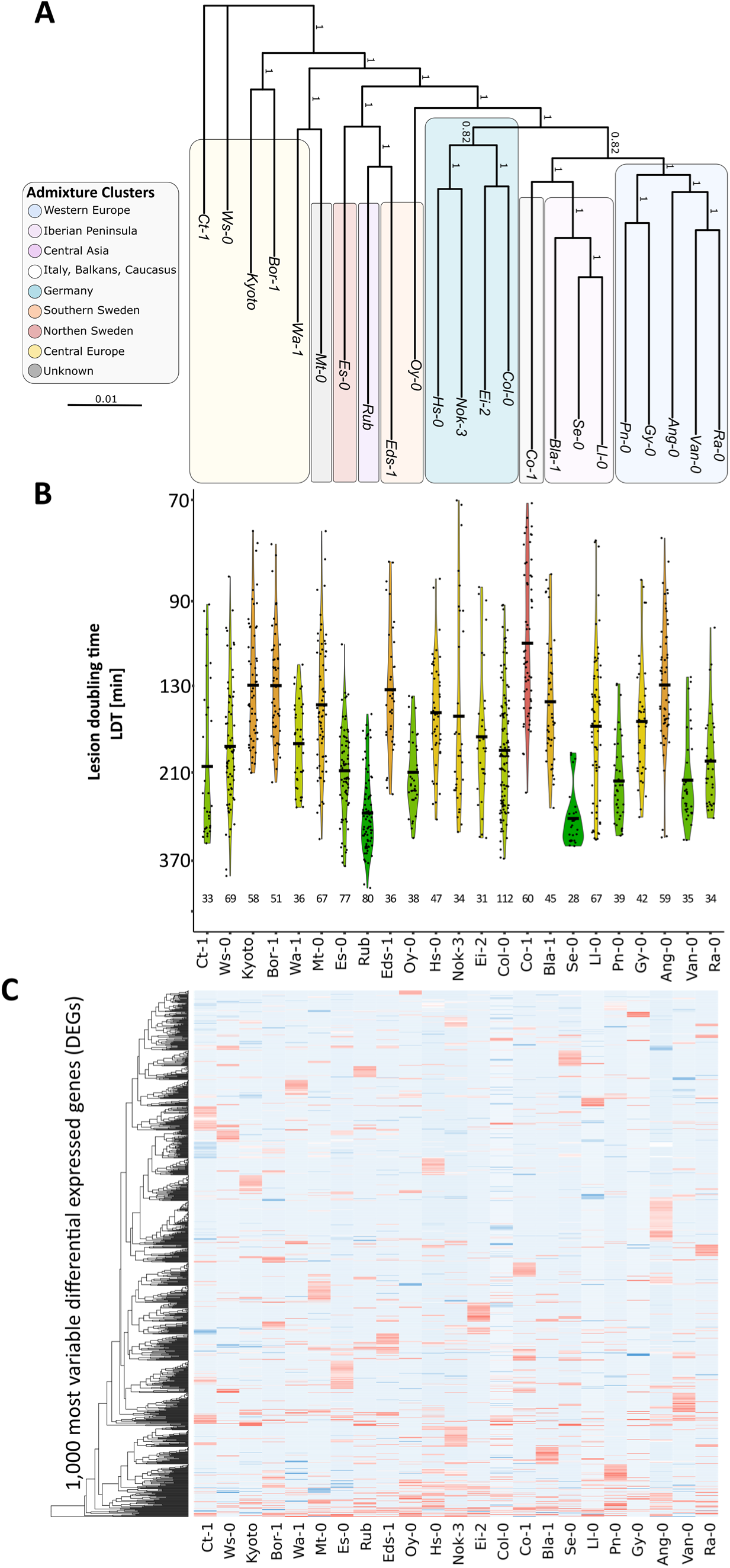
Global differential gene expression patterns and phylogeny do not correlate with *S. sclerotiorum* resistance across *A. thaliana* accessions. **A.** Phylogenetic tree of the 23 accessions used in this work based on coding sequences. Rectangles indicate the eight main admixture groups. Numbers along branches represent confidence values based on the UFBoot2 bootstrap method. **B**. Lesion doubling time (LDT) in minutes representing susceptibility of accessions at 24 hours after inoculation with *S. sclerotiorum*. Measurement were obtained from n=28 to 112 leaves in three experiments, with the black horizontal bar representing the mean speed. Kernels are colored according to mean susceptibility, from the most resistant (green) accessions to the most susceptible (red). **C**. Heatmap displaying the distribution of the 1,000 differentially expressed genes (DEGs) upon *S. sclerotiorum* inoculation with the highest coefficient of variation across accessions. Accessions are ordered based on genetic proximity in all panels.

± 0.11, Table S4-5). DEGs represented an average 39.56% ± 2.80% of expressed genes in each accession. Overall, 17,340 genes corresponding to 52.36% of *A. thaliana* transcriptome were differentially expressed in at least in one accession. Comparison between phylogenetic tree of accessions and transcriptome similarity dendrogram did not show clear congruence (Figure S2). To estimate the size of *S. sclerotiorum*-responsive transcriptome at the species level, we performed random accession sampling and logarithmic regression fit. This predicted a maximum of 22,643 DEGs (68.4% of the predicted transcriptome) across the whole *A. thaliana* diversity with R²=0.947 (Figure S3). Across accessions, 51.5% DEGs showed a maximum LFC variation < 2, and 2.6 % had a maximum LFC variation > 10 (Figure S4), indicating that drastic changes of expression profiles across accessions are limited to a few genes (Figure 1C). To assess the diversity of transcriptional responses to *S. sclerotiorum* at the species level, we compared the expression profile of the 17,340 DEGs across 23 accessions (Figure 2). DEGs with a differential expression in one accession only (accessory) ranged from 36 to 292 per accession, making a total 2,202 accessory DEGs (12.7%) at the species level. There were 13,181 DEGs (76%) differential in two to 22 accessions (shell). We identified 1,957 DEGs (11.3%) differential in all accessions (core) including 1,049 upregulated core DEGs and 908 downregulated core DEGs. Among the 17,340 DEGs, 1221 (7.0%) showed opposite regulation patterns (up-versus down-regulated) in at least two accessions. The number of DEGs with opposite regulation decreased from 13% for DEGs differential in two accessions to 0% in core DEGs (Figure 2).

**Figure 2:**
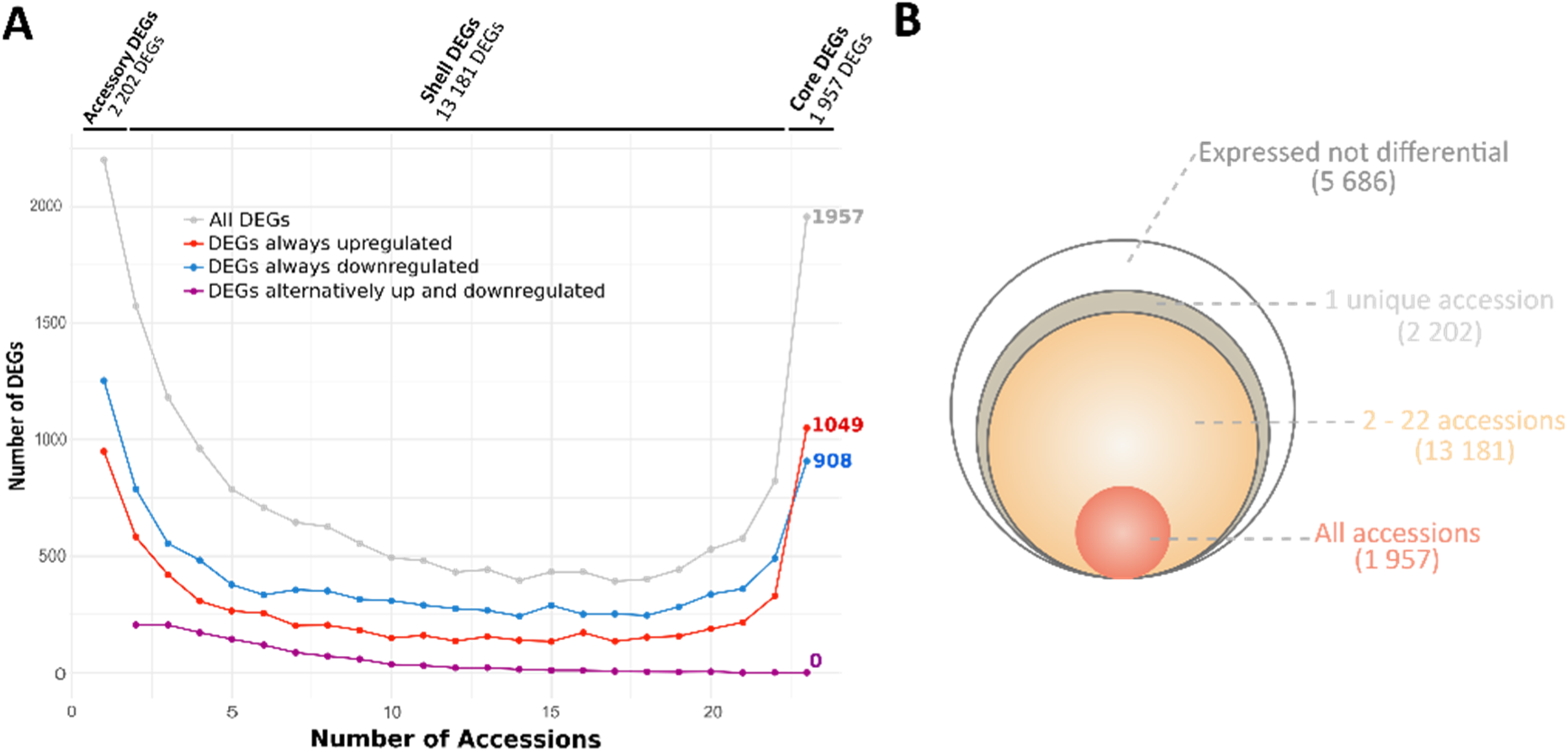
A total of 17,340 genes are differentially expressed in at least one accession. **A.** Number of differentially expressed genes (DEGs) according to the number of accessions in which they are differential. The graph shows DEGs upregulated in all accessions where they are differential (red), DEGs downregulated in all accessions where they are differential (blue), DEGs upregulated in at least one accession and downregulated in at least one accession (purple) and the total number of DEGs (grey). **B.** Synthetic representation of DEG classes, with the number of DEGs per class between brackets.

These results show that *S. sclerotiorum* infection triggers reprogramming of a majority of *A. thaliana* species pan-transcriptome, with moderate expression diversity across accessions and no clear link between the phylogenetic proximity of accessions and their transcriptomic response to *S. sclerotiorum*. This also emphasizes that diverse transcriptomes map to similar QDR phenotypes such as in the case of Rub and Se-0 (most resistant accessions) or Co-1 and Ang-0 (most susceptible accessions).

### A small set of strongly responsive genes defines *A. thaliana* core response to *S. sclerotiorum*

To characterize QDR responses conserved at the species level, we examined the expression, functional annotation and conservation of core DEGs. Core DEGs were consistently up- or down-regulated in all 23 accessions and showed a high expression stability, with 79% of core DEGs having an LFC standard deviation below 2. (Figure 3A). Up/down regulated DEGs ratio was 1.16 in core DEGs, 0.76 in accessory DEGs and 0.60 in shell DEGs, suggesting the accumulation of genes upregulated upon *S. sclerotiorum* inoculation during *A. thaliana* evolution. Using regression analysis, we estimated a minimum core DEG set of 719 upregulated and 352 down regulated genes across the whole *A. thaliana* diversity (Figure S5). The mean LFC values per gene show that core up- and down-regulated DEGs are among the most highly regulated genes in response to *S. sclerotiorum*. Of the top 500 DEGs with the highest mean LFC values, 72% are core up-regulated DEGs (6.54-fold enrichment compared to all up-regulated genes), while 45.5% of the top 500 down-regulated DEGs are core down-regulated (6.78-fold enrichment compared to all down-regulated genes) (Figure 3B). To determine the functional landscape of core DEGs, we performed a Gene Ontology (GO) enrichment analysis. GOs corresponding to 254 Biological Processes were significantly enriched among the upregulated core DEGs (Figure 3C). The majority of these GOs corresponded to ontologies related to primary metabolism, secondary metabolism, molecules transport and defense responses. Precisely, GOs related to secondary metabolism included camalexin biosynthetic process (GO:0010120), chorismate metabolic process (GO:0046417) and lignin biosynthetic process (GO:0009809), GOs related to defense responses included pattern recognition receptor signaling pathway (GO:0002221), response to ethylene (GO:0009723) and jasmonic acid mediated signaling pathway (GO:0009867). Core Up DEGs included known defense related genes such as *ABCG40*, *CERK1*, *PAD3*, *RBOHD* and *RLP30* (Stotz *et al*., 2011; Zhang *et al*., 2013; Zhou *et al*., 2013; Sucher *et al*., 2020; Pi *et al*., 2023). GOs corresponding to 154 Biological Processes were significantly enriched among the down regulated core DEGs (Figure 3D), including responses to light, developmental responses or photosynthesis. To study the evolutionary dynamics of core DEGs, we performed a phylostratigraphic analysis using Phytozome orthogroups. 1,039 upregulated core DEGs and 899 downregulated core DEGs were included in Phytozome orthogroups. Core *S. sclerotiorum*-responsive genes showed a clear gene age pattern with dominant old genes (Viridiplantae and Embryophyta) and young genes (Brassicaceae and *A. thaliana*) to a lesser extent, creating skewed U-shaped distributions (Figure 3E). These distributions are similar to those of all *A.thaliana* genes, indicating that *A. thaliana* transcriptome is predominantly composed ofby old and young genes. Compared to all *A. thaliana* genes, core DEGs included significantly fewer young genes and more old genes (+17.6% for upregulated core DEGs, +21.6% for downregulated core DEGs). Relative to *A. thaliana* whole genome, accessory DEGs included fewer old genes (−1.7%) while shell DEGs were enriched in old genes (+ 4.2%). These patterns suggest that intra-specific expression variation is better tolerated in young genes. To explore the relationship between transcriptome conservation and genetic diversity, we calculated nucleotide diversity (π and θ) in the 23 *A. thaliana* accessions. Core and shell DEGs exhibited significantly lower θ nucleotide diversity (−18.8% and -9.8% respectively) compared to genes not differentially expressed during infection (Figure 3F). The median non-synonymous to synonymous nucleotide diversity ratio (π_N_/π_S_ ratio) was 0.78 in core DEGs, 0.96 in shell DEGs, 1.05 in accessory DEGs and 1.22 in non-DEGs, indicating that most core DEGs are under purifying selection (Figure 3G). Therefore, conservation of gene expression patterns is linked to selective forces reducing nucleotide diversity and non-synonymous mutations.

**Figure 3.**
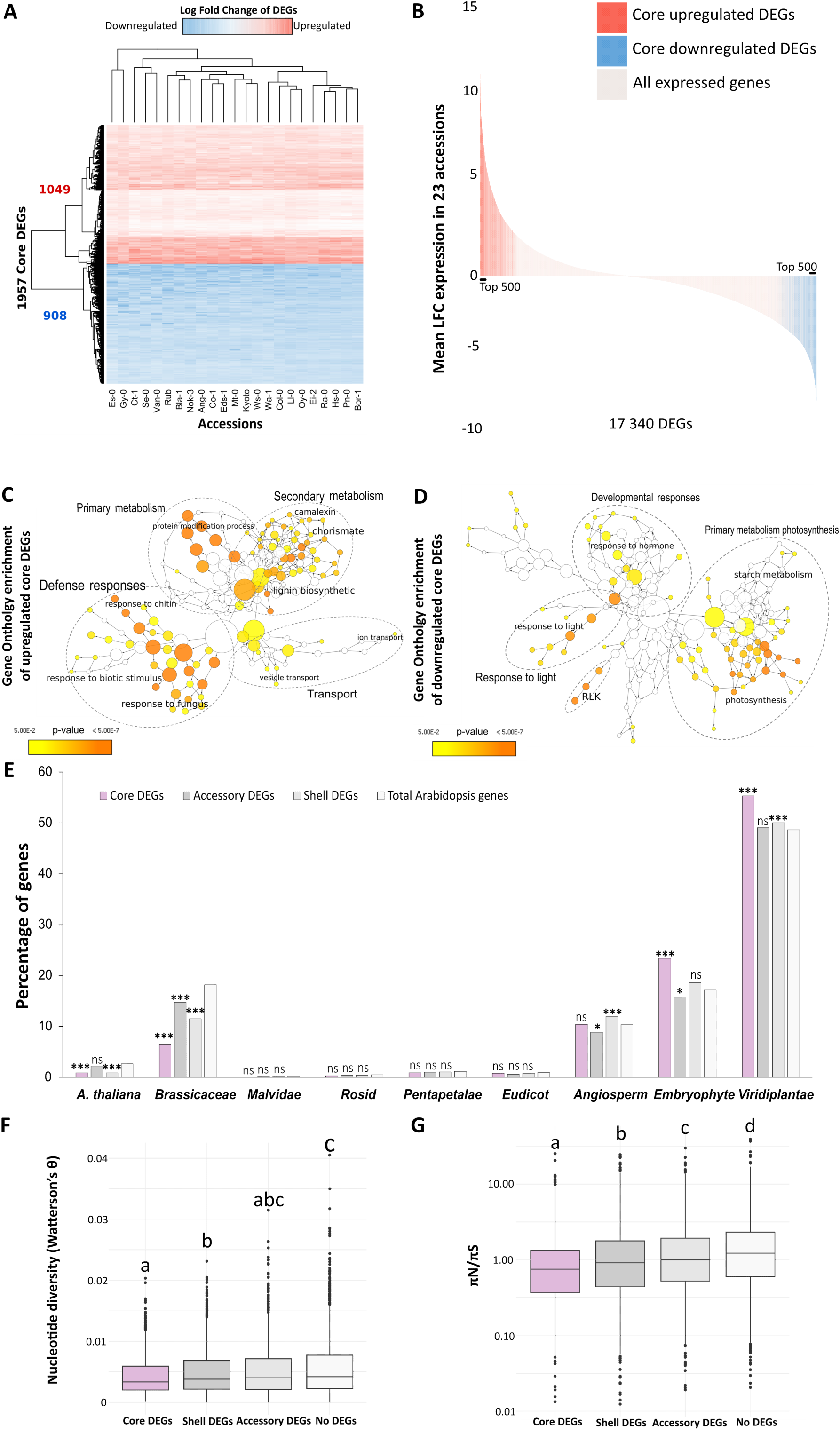
1.957 core DEGs are differentially expressed in all 23 *A. thaliana* accessions upon *S. sclerotiorum* inoculation. **A.** Heatmap displaying the distribution of 1.957 core differentially expressed genes (DEGs) across the 23 accessions after *Sclerotinia* inoculation, with an absolute log-fold change (|LFC|) ≥ 2. Red panels correspond to 1.049 genes significantly upregulated, and blue panels correspond to 908 downregulated genes. **B**. Mean LFC by gene across the 23 accessions. Red bars correspond to 1.049 core upregulated DEGs, and blue panels correspond to 908 core downregulated DEGs. **C and D.** Gene Ontology (GO) enrichment analysis of the 1.049 core upregulated DEGs (C) and the 908 core downregulated DEGs (D), plotted using the BINGO module from the Cytoscape software. Enrichment was calculated using BINGO and visualized on the GO hierarchy using Cytoscape. Circle size represents the number of genes in one GO category. **E.** Proportion of DEGs according to gene age. Gene ancestry information was retrieved from the Phytozome database. Red corresponds to core DEGs (upregulated and downregulated), grey to accessory DEGs, light grey to shell DEGs and white to the total genes in the Arabidopsis Phytozome database. Statistical analyses were conducted using the chi-squared test. Asterisks indicate the following p-values: * p < 0.05, ** p < 0.01, *** p < 0.001, and ns = not significant. **F and G.** Genetic population metrics in core DEGs (orange), shell DEGs (light blue) and accessory DEGs (blue). (F) Nucleotide diversity (θ) and (G) the non-synonymous to synonymous nucleotide diversity ratio (πN/πS ratio), with the y-axis scale plotted logarithmically. Letters identify significantly different gene groups in 1,000 bootstrap replicates of randomly sampled subsets of the same size using Dunn tests. One representative sample is shown in F and G.

We conclude that purifying selection at the species level drove the long-term sequence and function conservation, high and robust expression of *A. thaliana* core response genes to *S. sclerotiorum*.

### Transcriptome-wide co-expression clustering reveals the rewiring of gene modules across accessions

To compare the diversity of transcriptomic responses to *S. sclerotiorum* between accessions, we used log fold change (LFC) values for all expressed genes to correlate transcriptomes across accessions. Pearson correlation was calculated between each transcriptome to assess the overall proximity between accessions. Based on the similarity of global gene expression patterns, we identified subsets of accessions that were largely consistent with the DEG-based dendrogram (Figure 4A, Figure S2). We found no evidence for a link between correlation matrix clustering and susceptibility to *S. sclerotiorum*, or with environmental parameters at the geographic origin of the accessions (Table S6). Additionally, we observed no significant correlations between transcriptome subsets and phylogenetic clusters, indicating the absence of a phylogenetic signal in defining transcriptome proximity among accessions.

**Figure 4.**
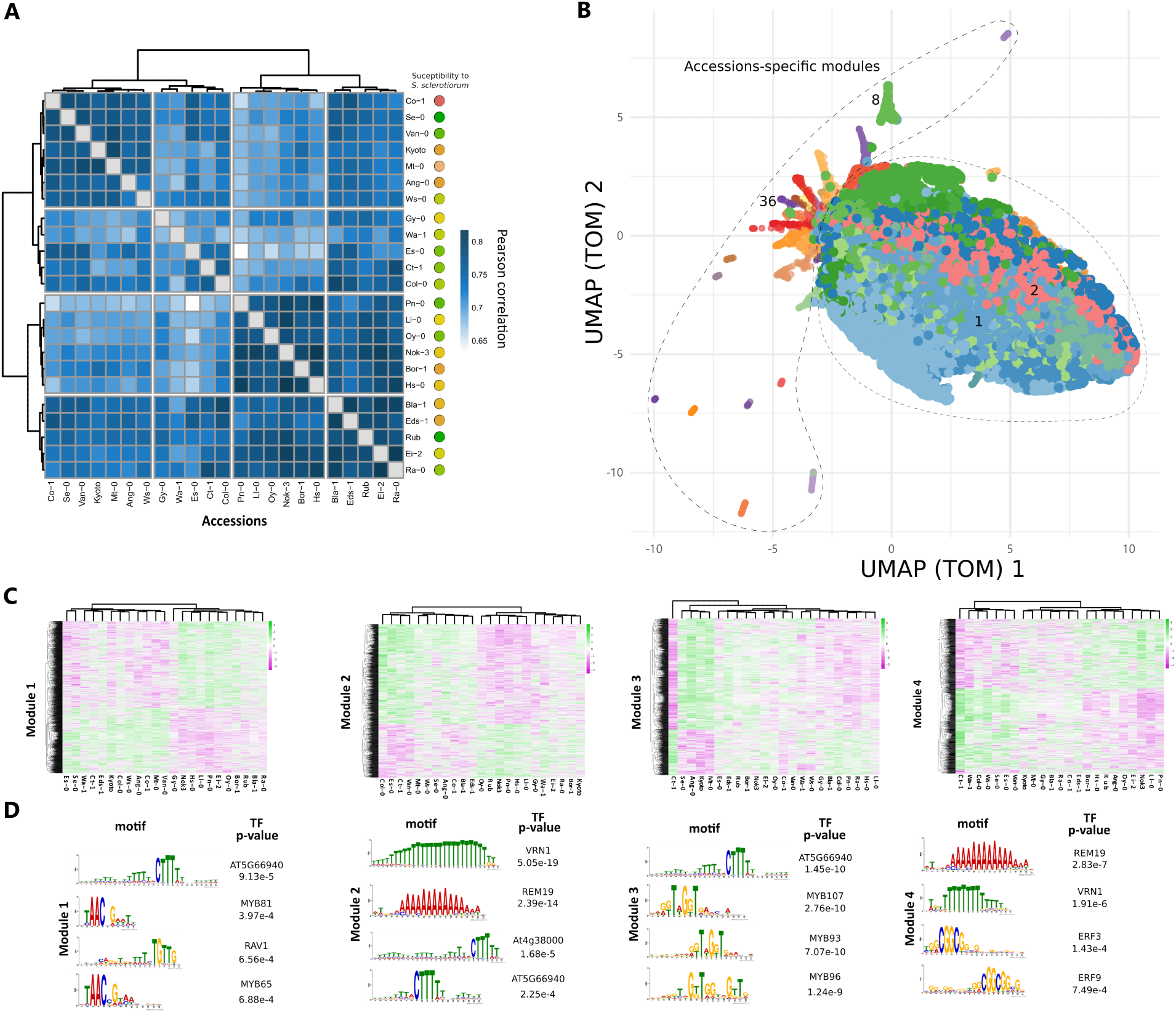
Identification of 48 gene co-expression modules in the response of *A. thaliana* accessions to inoculation by *S. sclerotiorum*. **A.** Pearson correlation matrix of the 23 accessions based on global transcriptome profiles upon *S. sclerotiorum* inoculation. Color circles indicate susceptibility to *S. sclerotiorum* for the 23 accessions according to Figure 1B. **B**. UMAP of TOM matrix obtained with the WGCNA co-expression analysis. Each color represents a distinct co-expression module. The dotted lines illustrate the accession-specific modules diverging from other gene co-expression modules. **C**. Heatmap showing the expression profile of genes associated with the four largest co-expression modules across 23 accessions. Green indicates LFC > 0, and purple indicates LFC < 0. **D**. DAP cis-motif enrichment comparing the 500 bp promoter regions in the two major accession groups identified by dendrogram clustering in genes from co-expression module 1 to 4. P-value corresponds to enrichment tests obtained with SEA from the MEME Suite, comparing the promoter genes from the smallest accession groups against the larger accession group described by the heatmap dendrogram in C, with TF corresponding to transcriptome binding of the DAP-seq motif. UMAP: Uniform Manifold Approximation and Projection; TOM: Topological Overlap Matrix; TF: Transcription Factors.

To further investigate transcriptome similarity in response to *S. sclerotiorum* across accessions, we performed transcriptome-wide co-expression module detection. Weighted correlation network analysis (WGCNA) identified 48 co-expression modules, comprising between 3 and 2,826 genes each, with a median of 314.15 genes per module (Figure 4B). Based on accession clustering, 25 co-expression modules were predominantly influenced by gene expression changes across multiple accessions, whereas 23 modules were driven by major gene expression changes specific to a single accession (Figure S6). Transcriptome similarity between accessions was not consistent across gene modules as illustrated by gene expression patterns from the four major modules (Figure 4C-F). For instance, the Spearman correlation between the accession dendrogram for module 1 and modules 2, 3 and 4 was weak, with values of 0.22, 0.04, and 0.33, respectively (Figure S7). GO enrichment analysis in the four major modules identified GOs related to response to stress in module 1, hormone signaling and primary metabolism in module 2, development and gene expression regulation in module 3, and immune responses in module 4 (Figure S8). This highlights the combination of diverse processes with contrasted quantitative contribution to the QDR response of each accession.

Enrichment in gene expression regulation ontologies in module 3 suggests that variation in cis may contribute to distinct QDR responses across *A. thaliana* accessions. To get support for cis-regulation variation into the diversity of *A. thaliana* QDR responses, we analyzed the distribution of DAP cis-motifs in the promoter of genes from each module. We compared motifs found in the promoter of accessions forming the two major similarity groups for each module. DNA binding sites for the transcription factors AT5G66940, MYB81, RAV1, and MYB65 were differentially enriched in module 1 genes (Figures 4G-H). A few motifs were differentially enriched in genes from several modules, indicated no strong signal of common DNA binding site variation across co-expression modules. But some motifs like MYB81, MYB96 and ERF3 motifs were uniquely enriched in genes from module 1, 3 and 4 respectively.

We conclude that regulatory variation targeting modules of shell DEGs, partly driven by cis-motif divergence between accessions, underlie diversity of the QDR transcriptome at the species level.

### Correlation between gene expression and disease susceptibility is marginal at the species level

To relate gene expression with disease susceptibility at the species level, we tested for correlation between gene expression and accessions susceptibility. First, we tested the correspondence between distance matrices for accession phenotypes and accession transcriptome similarity for each of the co-expression modules identified previously. Seven co-expression modules significantly explained the variation in resistance at the species level: five exhibited a positive correlation (Modules 38, 25, 26, 4, and 8) and three showed a negative correlation (Modules 20, 31, and 37) (Figure 5A). However, significant correlation values only ranged from 0.15 to 0.21 for positively correlated modules and -0.12 to -0.15 for negatively correlated modules. Except for co-expression modules 4 and 25, these correlations were driven by gene expression changes in a single accession (Nok3, Se-0, Wa-1, Ws-0, or Oy-0) (Figure S6). Therefore, variations in disease susceptibility at the species relate to the regulation of co-expression gene modules only in a limited number of accessions.

**Figure 5:**
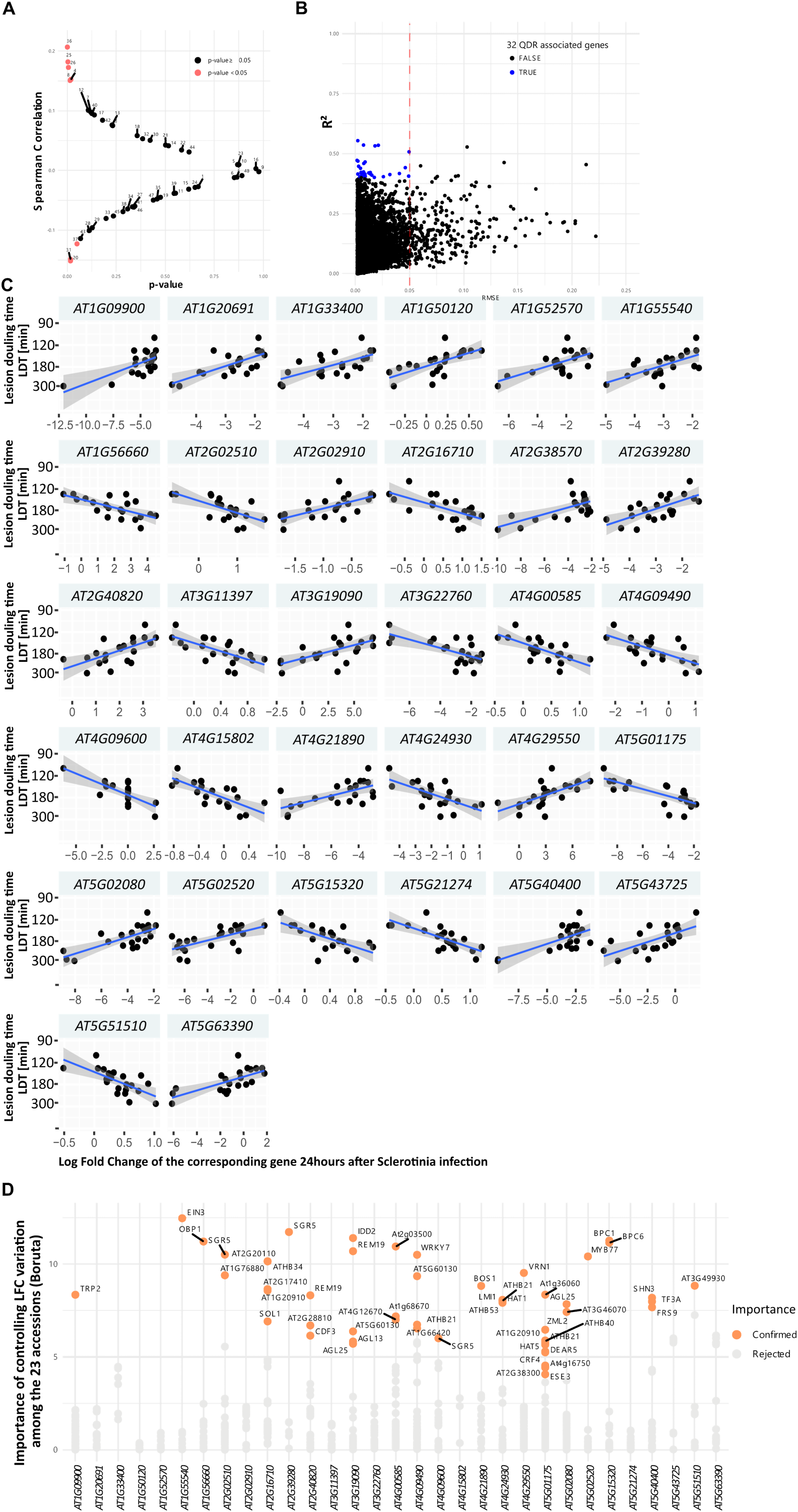
Analysis of species-wide correlation between gene expression and disease resistance phenotypes. **A.** Spearman correlation between dendrogram-derived distance matrices from gene expression clustering and phenotypic distance matrices for co-expression modules identified in Figure 3. P-values were calculated using a Spearman permutation test (n = 9,999). Co-expression modules with p-values < 0.05 are highlighted in red. **B**. Correlation between expression profiles obtained by RNA-seq and disease resistance in 23 accessions for all expressed genes. Blue dots represent genes with |R²| > 0.4 and RMSE < 0.05. The red dotted line indicates RMSE = 0.05. **C**. Correlation between LFC (X-axis) and disease resistance (Y-axis) in 23 accessions for the 32 best correlated genes. For each gene, the 21 out of 23 log fold change (LFC) values with the lowest p-values were retained. Genes with |R²| > 0.4 and RMSE < 0.05 are presented in the figure. The blue lines show linear regression of the data with R² and p-value labelled and the 95% confidence interval shown as a grey area. **D**. DAP cis-regulatory motifs with importance scores identified by the Boruta package as significant contributors to LFC variation across 23 accessions in the 32 best correlated genes.

Next, we tested if the expression of individual genes associated with QDR across accessions. For this, we used expression LFC for all expressed genes and disease resistance phenotype for correlation and regression analysis. With a |R²| > 0.4 and rmse < 0.05, the expression of 32 genes was significantly correlated with resistance variation, none of which had been previously associated with disease resistance to *S. sclerotiorum* (Figure 5B-C). Among those, 22 genes had an expression correlated with disease resistance (ie. R² < 0 and LFC > 0 or R² > 0 and LFC < 0), such as genes encoding the TPR9 tetratricopeptide repeat protein and the AT1G56660 MAEBL protein (Figure 5C). 10 genes had an expression correlated with disease susceptibility (ie. R² > 0 and LFC > 0 or R² < 0 and LFC < 0), including gene encoding the AT2G40820 Actin binding-like protein and the AT5G01175 microtubule controlling protein (Figure 5C). Among the 32 QDR-associated genes, 22 were shell DEGs differentially expressed in 2 to 22 accessions with an average |LFC| from 0.51 to 5.31. Ten genes had |LFC| ≤ 2 in all 23 accessions. This approach suggests that only a few shell DEGs were associated with QDR variation at the species level, potentially due to the limited contribution of individual genes or complex genetic interactions.

To investigate the regulatory mechanisms driving transcriptome variation at the species level and the acquisition of expression profiles of the 32 QDR-associated genes, we conducted a comparative analysis of the cis-motif content within the 500 bp promoter regions of these genes across 23 accessions. We identified 497 cis-motifs that showed at least one presence/absence polymorphism in the 500 bp promoter region the 32 QDR-associated genes. Next we used a random forest approach to identify a set of 46 cis-motifs whose presence/absence polymorphism in promoter regions robustly associated with LFC variation in at least one of the 32 genes (Figure 5D, Table S7). Specifically, the ATHB21 and SGR5 cis-motifs were responsible for LFC variation in three genes (*AT4G09490*, *AT4G24930* and *AT4G24930* or *AT2G02510*, *AT2G39280* and *AT4G09600* respectively), the four cis-motifs AGL25, AT1G20910, AT5G60130, and REM19 were responsible for LFC variation in 2 genes (Table S7), while 40 cis-motifs influenced LFC variation in a single gene. Notably, the impact of these cis-motifs varied between genes, indicating a lack of consensus in cis-motif acquisition at the species level.

### Core genes make the transcriptome-phenotype map of *A. thaliana* QDR navigable

To get insights into how global gene expression constrains the evolution of QDR at the species level, we constructed a transcriptome-phenotype map of *A. thaliana* QDR against *S. sclerotiorum*. For this, we clustered the 23 accessions based on the log fold change (LFC) of all expressed genes using principal component analysis (PCA), and used Gaussian interpolation to connect experimentally-determined disease resistance phenotypes by a smooth surface (Figure 6A, B). This map describes the expected QDR phenotype for every point of the transcriptome space covered by our sampling of accessions. It is composed of two major sectors: a susceptibility basin connecting accessions Co-1, Ang-0, Mt-0, Kyoto, Eds-1, Gy-0, Bla-1, Bor-1, Nok-3 and Hs-0, and a resistance plateau connecting accessions Es-0, Ct-1, Van-0, Col-0, Ws-0, Wa-1, Ra-1, Ei-2, Rub, Oy-0 and Ll-0. Accessions Se-0 and Pn-0 form two resistance peaks at two sides of the map. Within this map, accessions distant in transcriptome space can be connected through similar phenotypes. This defines the equivalent of neutral networks of accessions connecting diverse transcriptomes with a similar fitness, approximated here by resistance to *S. sclerotiorum*.

**Figure 6:**
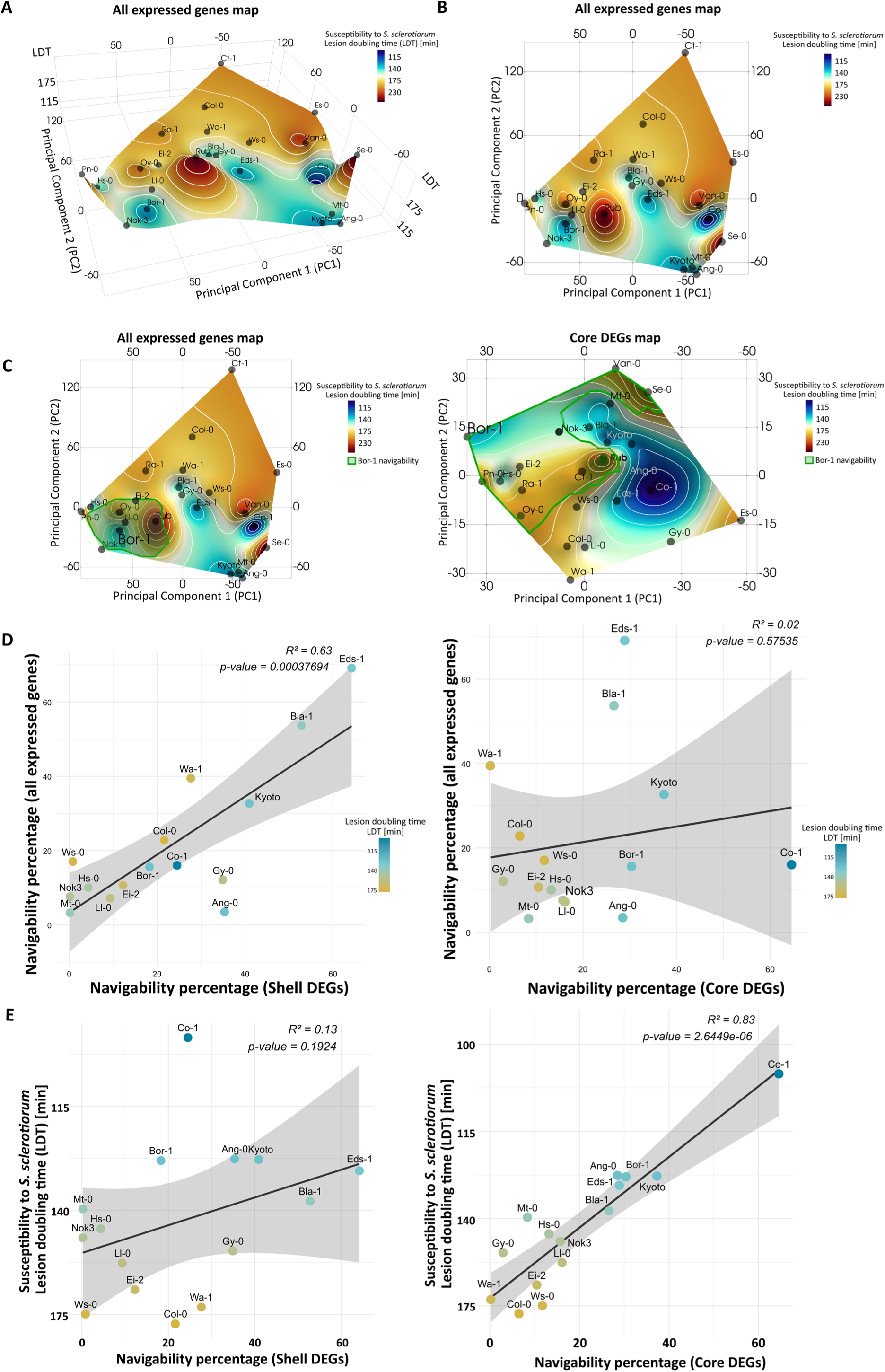
Navigability within the transcriptome - resistance map based only on core DEGs is correlated with susceptibility. **A.** Transcriptome-resistance map showing accession proximity in the transcriptome space in response to *S. sclerotiorum* inoculation (X-Y plane) and resistance to *S. sclerotiorum* (Z-axis, color scale). The x and y axes represent components 1 and 2 of a principal component analysis using LFC data for all expressed genes. The color gradient represents disease resistance from more susceptible (blue) to more resistant (red), with experimentally studied accessions shown as small labelled spheres. Lines of equal resistance (white) are provided for legibility **B.** Top view of the transcriptome-resistance map shown in A. **C** Top projections of the transcriptome-phenotype map for all expressed genes (left) and core DEGs only (right) showing sectors of the map accessible to Bor-1 accession without loss of resistance (navigability) delineated by green borders. **D** Relationship between navigability of the transcriptome-resistance map obtained for all expressed genes (Y-axis) and navigability of the map obtained for shell DEGs (X-axis, left) and core DEGs (X-axis, right) in 15 *A. thaliana* accessions. **E**. Relationship between resistance against *S. sclerotiorum* (Y-axis) and navigability of the transcriptome-resistance map obtained with shell DEGs (X-axis, left) and core DEGs (X-axis, right) in 15 *A. thaliana* accessions. The black lines show linear regression of the data with R² and p-value labelled and the 95% confidence interval shown as a grey area.

Genotype variants connected only through fitness increase define the navigability of fitness landscapes. We calculated the QDR landscape navigability for each accession, defined as the percentage of the transcriptome map accessible with no loss of resistance. Navigability values ranged from 69.14% (Eds-1) to 0.02% (Se-0) (Table S8). The resistance maxima observed in our study for the Se-0 and Rub genotypes can be reached by 14 genotypes without passing through susceptible valleys, due to the large intermediate resistance plateau (Figure 6B). This suggests that transcriptomic responses to *S. sclerotiorum* are highly evolvable in *A. thaliana*, tolerating significant changes with no resistance loss.

To evaluate the contribution of core and shell DEGs to *A. thaliana* QDR evolvability, we generated transcriptome-phenotype maps based on the expression of core DEGs only and all expressed genes map. We calculated the QDR landscape navigability for each accession on the core DEGs map and all expressed genes map (Table S8). QDR landscape navigability increased in 14 accessions when using transcriptome-phenotype maps based solely on the expression of core DEGs, compared to maps using all expressed genes. For example, the Bor-1 accession shows a navigability of 30.4% with the core DEGs-only map, compared to 15.6% with the all-expressed-genes map (Figure 6C, Figure S9). Navigability on the whole transcriptome map correlated with navigability on the shell DEG map (R² = 0.63, p-value = 0.00038) but not with the navigability on the core DEG map (R² = 0.02, p-value = 0.58), consistent with the prevalent contribution of shell DEGs to the overall response to *S. sclerotiorum* (Figure 6D). Susceptibility to *S. sclerotiorum* correlated with navigability on the core DEG map (R² = 0.83, p-value = 2.6e-6) but not with the navigability on the whole transcriptome map (R² = 0.01, p-value = 0.77) and the shell DEG map (R² = 0.13, p-value = 0.19) (Figure 6E) indicating that QDR evolvability is related to regulatory variation in core DEGs in this fitness landscape.

## Discussion

Quantitative disease resistance provides partial and durable protection against various pathogenic microbes across the green lineage. Genetic mechanisms associated with the short and long-term evolution of QDR remain largely obscure (Kou and Wang, 2010). In a previous study, comparative transcriptome analysis across plant species highlighted exaptation through regulatory divergence as a path towards the evolution of QDR against *S. sclerotiorum* (Sucher *et al*., 2020). Here we analyzed patterns of QDR transcriptome evolution at the species level through a comprehensive RNAseq analysis of 23 *Arabidopsis thaliana* accessions displaying various degrees of resistance to *S. sclerotiorum*.

### How complex is plant quantitative disease resistance?

The omnigenic model of complex traits proposes that most genes expressed in an infection context should contribute to the heritability of disease resistance, either through a role in immunity or in its regulation (Boyle *et al*., 2017). Interspecific comparison found an average 32.63% genes responsive to *S. sclerotiorum* across Pentapetalae, an estimate resulting from the analysis of a single genotype of each species (Sucher *et al*., 2020). Here we assessed infra-specific variation in *Arabidopsis thaliana* gene responsiveness to *S. sclerotiorum*. In each accession, DEGs represented an average 39. 6% of expressed genes, together accounting for 52.4% of *A. thaliana* pan-transcriptome responsive to *S. sclerotiorum* infection. This number is consistent for instance with 50.4% of *A. thaliana* genes differentially regulated upon *Fusarium oxysporum* inoculation (Guo *et al*., 2021) and suggests that infra-specific variation may expand the repertoire of QDR genes by ∼1.3 fold. We have chosen to map RNA-seq reads to the TAIR10 reference genome for a more accurate quantification of gene expression. This approach does not allow to identify accession-specific transcripts that are absent from the reference genome, and probably slightly underestimates the overall diversity of *S. sclerotiorum* responsive transcripts in *A. thaliana*.

### How predictable is quantitative disease resistance?

There was no detectable phylogenetic signal associated with global transcriptome patterns and resistance phenotype in our 23 *A. thaliana* accession panel, indicating that the heritability of these traits at the species level is complex. Previous analyses reported that transcriptome variation tends to recapitulate evolutionary relationships accurately at the interspecies level but loses precision for closely-related species (Winkelmüller *et al*., 2021; Wu *et al*., 2022). This may be due to transcriptional noise being prevalent in a subset of QDR genes, or to the independent evolution of sequence and regulatory polymorphisms at QDR loci (Tirosh and Barkai, 2008; Harrison *et al*., 2012; Uebbing *et al*., 2016). Contrasted contributions of regulatory divergence and sequence variation to the evolution of quantitative plant immunity may indeed result in independence between these polymorphisms at the species level. QDR phenotypes may therefore be difficult to predict from sequence information alone (Gentzbittel *et al*., 2019). By training Machine Learning (ML) models on plant transcriptomes, Sia *et al*. found that information on expression of only 0.5% of genes was sufficient to predict classes of disease phenotypes across multiple pathosystems (Sia *et al*., 2023). Here, we identified 32 genes whose expression correlated with QDR phenotypes at the species level. They have not been associated with disease resistance to date but may encodes putative signaling proteins controlling known defense pathways, and could potentially be used as predictors of QDR phenotypes at the species level. However, the coordinated evolution of gene expression at the sub-species level could contribute to phenotypic adaptation (Mezey et al., 2008), and impede phenotypic predictions unless the genetic diversity of the species has been investigated with sufficient depth (Crossa *et al*., 2010).

### What is the contribution of the variable transcriptome to *A. thaliana* QDR?

Regulatory divergence across *A. thaliana* accessions was generally of low magnitude, questioning the adaptive value of this diversity. By contrast to tightly regulated core genes, shell and accessory genes identified in our work may exhibit a high level of transcriptional noise, possibly underlying a bet-hedging strategy to improve immunity (Viney and Reece, 2013; Urban and Johnston, 2018). In such a strategy, noise may lead to stochastic activation of diversified immune responses in cell subsets. This local immune response may be sufficient to limit pathogen colonization or may propagate through cell to cell communication. Over long evolutionary time, the plasticity of gene expression may also facilitate the acquisition of precise temporal and spatial expression patterns underlying QDR (Groen *et al*., 2020; Jones and Vandepoele, 2020). The recent advent of single-cell RNA-sequencing (Tang *et al*., 2023) will allow testing cell population immunity mechanisms that were not accessible to bulk RNA analyses. The existence of 32 genes whose expression correlated with QDR at the species level suggests that minor transcriptional variations in key regulators may play a significant role in phenotypic evolution due to their interactions within the broader gene networks (Liu *et al*., 2019; Jenull *et al*., 2021). Exploring regulatory mechanisms and functions of genes whose expression varies across species genes presents an opportunity to identify new contributors to QDR.

### What is the contribution of core DEGs to *A. thaliana* QDR phenotype?

We identified 1049 core up-regulated Among *S. sclerotiorum*-responsive genes in *A. thaliana* in all accessions, suggesting either a role in mediating resistance across various genetic backgrounds (Sucher *et al*., 2020; D., Yang *et al*., 2021) or broad-spectrum manipulation by the fungus. Gene Onthology analysis associated core upregulated DEGs with general responses to fungal pathogens, such as the biosynthesis of camalexin, chorismate metabolism, and lignin biosynthesis (Zhu *et al*., 2013; Zhang *et al*., 2017). These processes, conserved at the species level, typically contribute to a broad-spectrum defense against fungal pathogens, including *Botrytis cinerea*, *Colletotrichum higginsianum*, and *Phytophthora parasitica* (Berre *et al*., 2017; Nicole E. Soltis *et al*., 2020; Zhu *et al*., 2023). Core DEGs form a backbone network where fine-tunable transcriptome modules play a pivotal role in responding to pathogen genetic diversity (Kim *et al*., 2014; Hillmer *et al*., 2017). Mutants *pad3-1*, *abcg40-2* and *rlp30-1* altered in core DEGs show increased susceptibility to *S. sclerotiorum*, indicating clear contribution of some core DEGs to QDR (Zhang *et al*., 2013; Sucher *et al*., 2020; Kusch *et al*., 2022). However, the relative contribution of many core DEGs to disease resistance remains to be tested, and why the global pool of core DEGs contributes to different levels of resistance across accessions is unclear. The contribution of core genes to QDR may be regulated at the post-transcriptional level in a contrasted manner across accessions. Alternatively, specific core transcript ratio may be required for efficient QDR, or the precise timing of core transcripts activation may be crucial for QDR, emphasizing the dynamic role of transcriptional reprogramming as a major contributor to fungal resistance (Niks *et al*., 2015; French *et al*., 2016; D., Yang *et al*., 2021).

### How robust and evolvable is QDR in *A. thaliana*?

Our analysis emphasizes redundancy in the transcriptome-phenotype map, where multiple global transcriptomes map onto the same level of disease resistance to *S. sclerotiorum*, defining neutral networks (Greenbury *et al.,* 2022). Neutral networks intrinsically support phenotypic robustness to (epi)genetic interference. Our fitness landscape model predicts that 15 accessions can navigate >1% of the transcriptome space with no loss of resistance. Most of the accessions tested had intermediate resistance allowing strong navigability, in agreement with QDR being predominant in natural populations (Corwin and Kliebenstein, 2017). High transcriptome navigability could be a crucial component for the durability of QDR, by preventing extreme fitness valleys or peaks that limit transcriptome flexibility. A corollary of this finding is that globally similar transcriptomes can produce very contrasted disease phenotypes (One Thousand Plant Transcriptomes Initiative, 2019). This may be explained by a prevalent role of regulation at the post-transcriptional level in QDR, although the massive transcriptional reprogramming induced by *S. sclerotiorum* inoculation argues against this hypothesis. Alternatively, the QDR response network may be conserved and redundant, or structured to canalize similar downstream responses in spite of variable inputs (different pathogen genotypes, environmental conditions, or plant genetic background) (Zhang *et al*., 2017). Neutral networks also allow populations to access a wide variety of transcriptomes and novel phenotypes without intermediate fitness penalty, promoting evolvability (Greenbury *et al*., 2016). Broad neutral networks of QDR transcriptome support high evolvability of this trait at the species level. It should be noted that our focus on a limited sampling of 23 accessions may underestimate the full range of the navigable transcriptome space, or conversely have missed fitness valleys that limit navigability. Besides, our transcriptome-phenotype maps only represent the two principal components of the transcriptome space, and navigability may vary when considering additional dimensions of this space (Zagorski *et al*., 2016).

### How did the regulation of fungal-responsive genes evolve?

Variation in transcriptome may be facilitated by regulatory elements or epigenetic mechanisms that alter gene expression even before genetic variants arise in the population (Alvarez *et al*., 2015). The acquisition of a new expression pattern in a limited number of genes may then significantly alter QDR phenotypes (Chen *et al*., 2013). The cross-accession comparison of DAP motifs in the promoter of genes forming co-expression modules and genes whose expression correlated with QDR identified presence/absence polymorphisms associated with regulatory variation. This variation often targeted a few accessions only. Our analyses point towards the acquisition of WRKY binding motifs, known for their involvement in transcriptional regulation (Wani *et al*., 2021) as a potential mechanism for the stabilization of core gene expression at the species level. When comparing the promoter sequences of 32 QDR-associated gene between the 23 accessions, we identified DAP motifs whose presence or absence is significantly associated with 21 gene expression patterns, such as the *cis* DNA binding motif recognized by the transcription factor WRKY7 for the QDR-associated gene *AT4G09490*. Most of the variation in cis motifs is unique to each QDR-associated gene, suggesting a possible recent reconfiguration of neutral transcriptome by accessions through mutations in the promoter regions (van Kooten *et al*., 2021). Given the unique DAP motifs enriched specifically in QDR responsive genes of each accession, we propose that the gain or loss of DNA TF-binding motifs contributes significantly to transcriptional variation at the species level. Contrary to our expectations, we did not find transposable element families significantly associated with QDR transcriptome modulation at the species level. Cis-regulatory variations arise from the emergence of novel enhancer sequences (Long *et al*., 2016) or from genome rearrangements that recruit new genes to transcriptionally active regions. Cis-regulatory variations may also serve as a source for generating allele-specific transcripts involved in QDR (Fukuoka *et al*., 2014; P., Yang *et al*., 2021). Novel enhancer activities often result from the co-option of transcription factor binding sites already present in ancestral enhancers (Wittkopp and Kalay, 2012; Macquet *et al*., 2022). The acquisition of cis-regulatory enhancers in promoter regions is known to drive the evolution of gene expression, shaping transcriptome patterns related to plant resistance (Ogasahara *et al*., 2022). The divergence of cis-regulatory motifs is suggested to be a key contributor to the rapid transcriptional reprogramming essential for the broad-spectrum efficiency of plant immune responses (Mine *et al*., 2018). Consequently, variation in cis-regulatory elements across genotypes depends on various selective forces, including abiotic and biotic environmental factors (Ricci *et al*., 2019; Marand *et al*., 2023).

### Which properties of the QDR transcriptome space promote robustness and evolvability?

The accumulation of multiple and independent mutations in the promoter region may lead to the rewiring of a neutral transcriptome (Greenbury et al., 2022), supporting high evolvability dynamics. Subtle differences in evolvability could play an important role in shaping the long-term success of transcriptomic variants (Ferrare and Good, 2024). The navigability of our core DEG transcriptome-phenotype map correlated with accession susceptibility, suggesting that purifying selection stabilized core DEGs expression to serve as a network backbone for QDR phenotype and genetic architecture to evolve. On the other hand, lineage-specific genes often participate in adaptation to biotic and abiotic stress (Tian *et al*., 2012). Thus, genes from the most recent age classes may contribute relatively directly to QDR, including a few with expression directly correlated to the QDR phenotype. The evolution of robust QDR phenotypes at the species level may involve concerted regulatory variation between the stable and reduced set of core genes and the extensive and plastic pathogen-responsive transcriptome, thud indicating the need to study evolution of the defence network as a whole (Kahlon & Stam, 2021). Future research should evaluate the impacts of environmental constraints on these evolutionary dynamics to explore both past and future trajectories of QDR evolution in changing environments.

## Acknowledgements

We thank Sébastien Carrère and the LIPME bioinformatics platform for assistance with data storage and manipulation including reads mapping. We are grateful to Marielle Barascud and Rémy Vincent for assistance in sample collection for RNA sequencing, and to the QIP group at LIPME for discussions and suggestions. This work was supported by the French Laboratory of Excellence project ’TULIP’ (ANR-10-LABX-41; ANR-11-IDEX-0002-02), l’Agence Nationale pour la Recherche (ANR-19-CE20-15, ANR-21-CE20-30), the European Research Council (ERC-StG-336808), and the INRAE SPE division.

## Conflict of interest statement

The authors declare that they have no competing interests to disclose.

## Notes

### Competing Interest Statement

The authors have declared no competing interest.

### Summary of Updates

The main updates are summarized as follows: 1) Improved transcriptome-phenotype association tests: We performed co-expression and correlation analyses across all 23 Arabidopsis thaliana accessions to identify QDR-associated genes. 2) Refinement of cis-regulatory region identification: Promoter sequences across the 23 accessions were analyzed to detect motifs linked to variations in QDR-related gene expression. 3) Conceptualization of QDR evolvability: We mapped transcriptome-phenotype relationships to explore evolutionary pathways that enhance QDR through transcriptomic changes. 4) Clarification of statistical methods: We thoroughly revised the explanation of controls, statistical analyses, and association tests for clarity and robustness.

