## Supplemental Figures S1-9 for "Neutral transcriptome rewiring promotes QDR evolvability at the species level"

### Supplementary Data

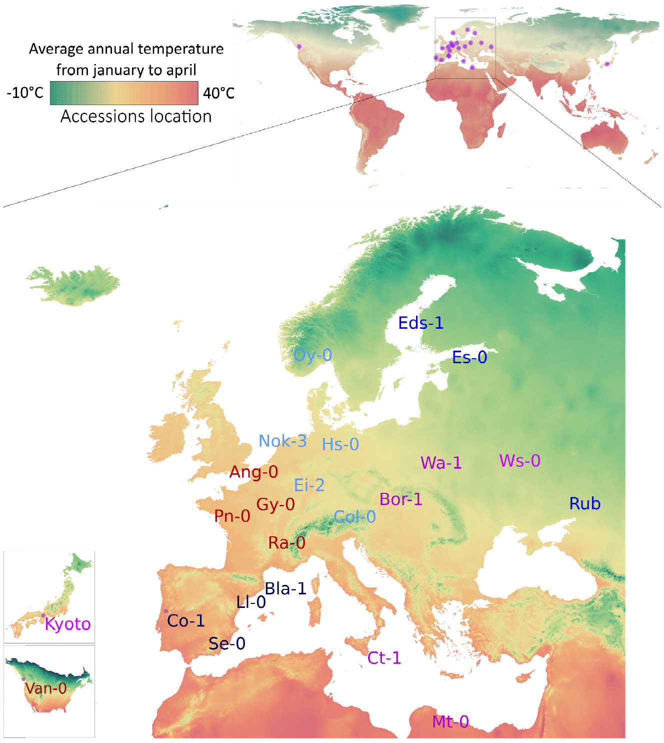


**Supplemental Figure S1: 23 natural accessions covering a broad range of the species geographic diversity.**
Distribution of the 23 accessions across Europe, Canada, and Japan. Purple dots represent the coordinates of the initial collection locations for each accession. The background color indicates the mean spring temperature, with blue indicating the coldest regions and red indicating the warmest.

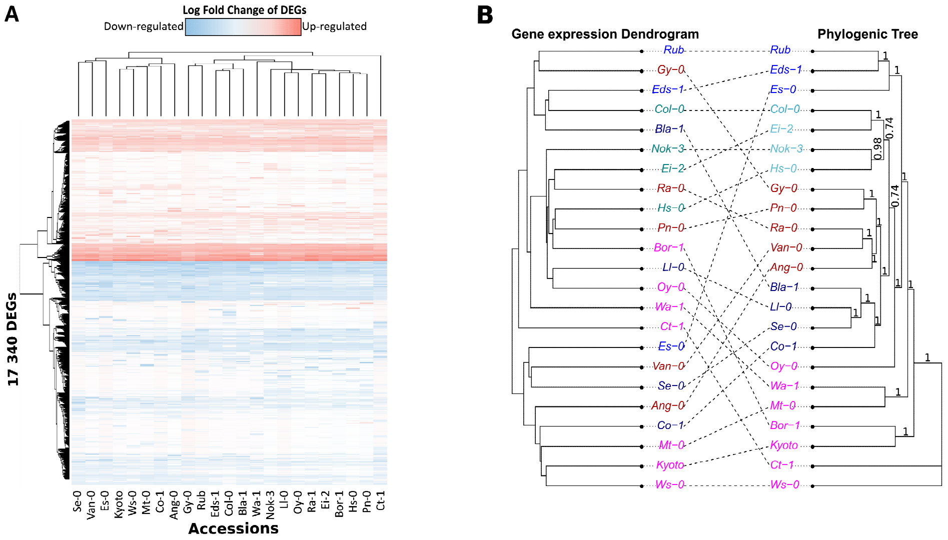


**Supplemental Figure S2: Phylogenetic tree of accessions and gene expression dendrogram did not show clear congruence.**

**A**. Heatmap displaying the distribution of 17.340 significantly differentially expressed genes (DEGs) in the 23 accessions after Sclerotinia infection, with |LFC| ≥ 2. **B**. Tanglegram comparing the gene expression dendrogram in Figure 1B to the phylogenetic tree based on coding sequences (CDS) across the 23 accessions. The label colors indicate the five main phylogenetic clusters, and the numbers at the branches represent confidence values obtained using the UFBoot2 bootstrap method.

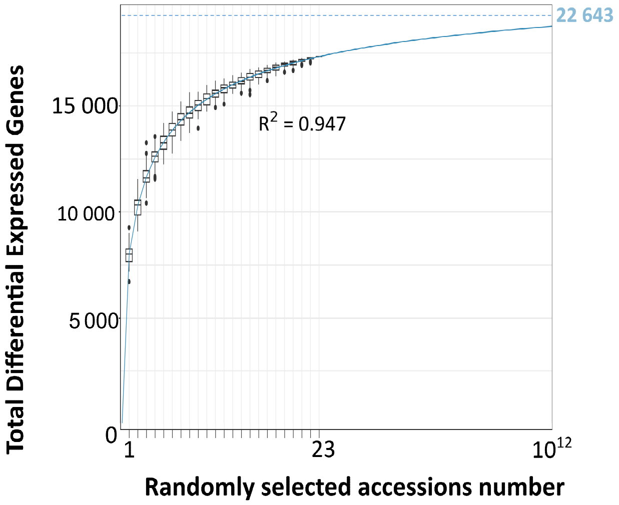


**Supplemental Figure S3: A maximum of 22,643 DEGs (68.4% of the predicted transcriptome) is estimated across the whole *A. thaliana* diversity.**
Regression analysis estimating the maximum number of DEGs through random sampling from 1 to 23 accessions. The regression curve, with an $R^2$ coefficient of 0.947, suggests a maximum of 22,643 DEGs with $10^{12}$ accessions.

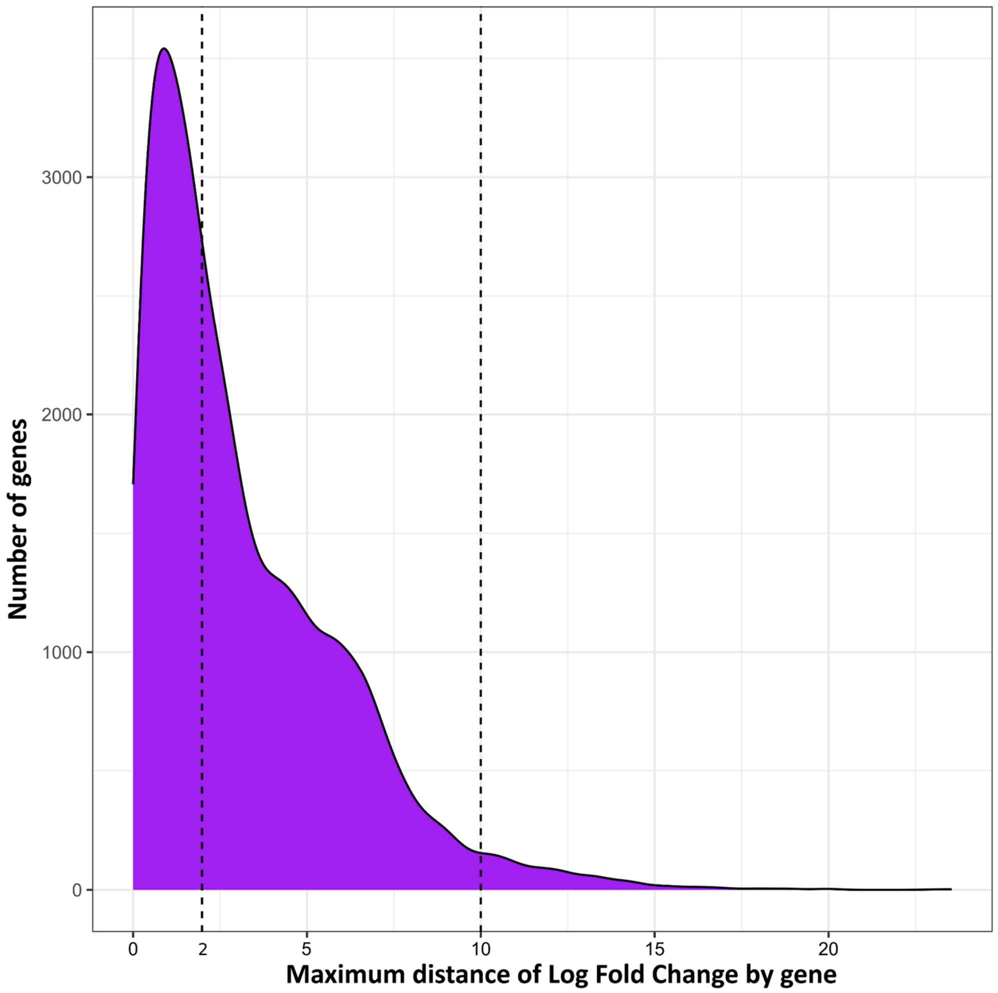


**Supplemental Figure S4: Maximum distance of Log Fold Change by gene.**
Density plot illustrating the maximum distance of Log Fold Change for each genes. The vertical dashed line indicates a maximum distance of 2 and 10 Log Fold Change.

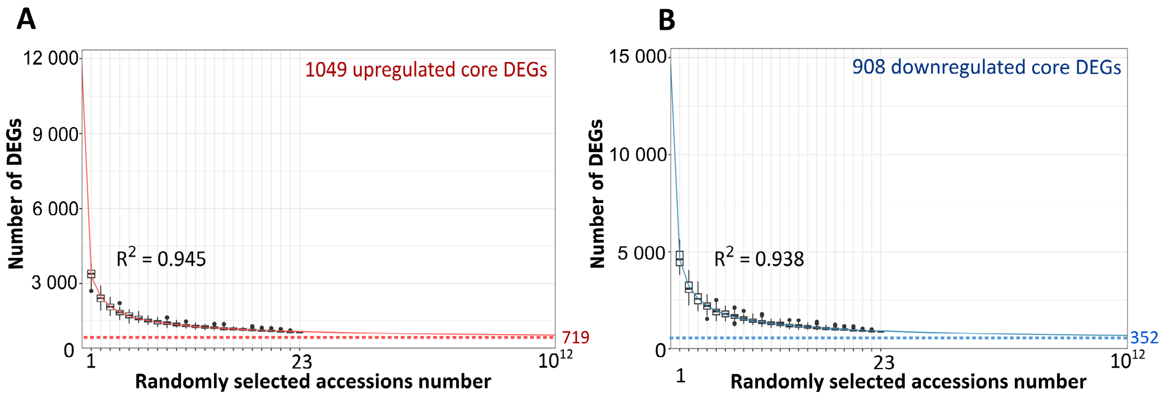


**Supplemental Figure S5: A maximum of 719 and 352 DEGs is estimated for respectively core upregulated DEGs and core downregulated DEGs considering the whole *A. thaliana* diversity.**

**A and B.** Exponential regression analysis estimating the minimal number of core upregulated (A) or downregulated (B) DEGs through random sampling from 1 to 23 accessions. The regression curves, with a regression coefficient of $R^2$= 0.945 and $R^2$= 0.938 respectively, suggest a minimum of 719 core upregulated DEGs and 352 core downregulated DEGs as the number of accessions is $10^{12}$.

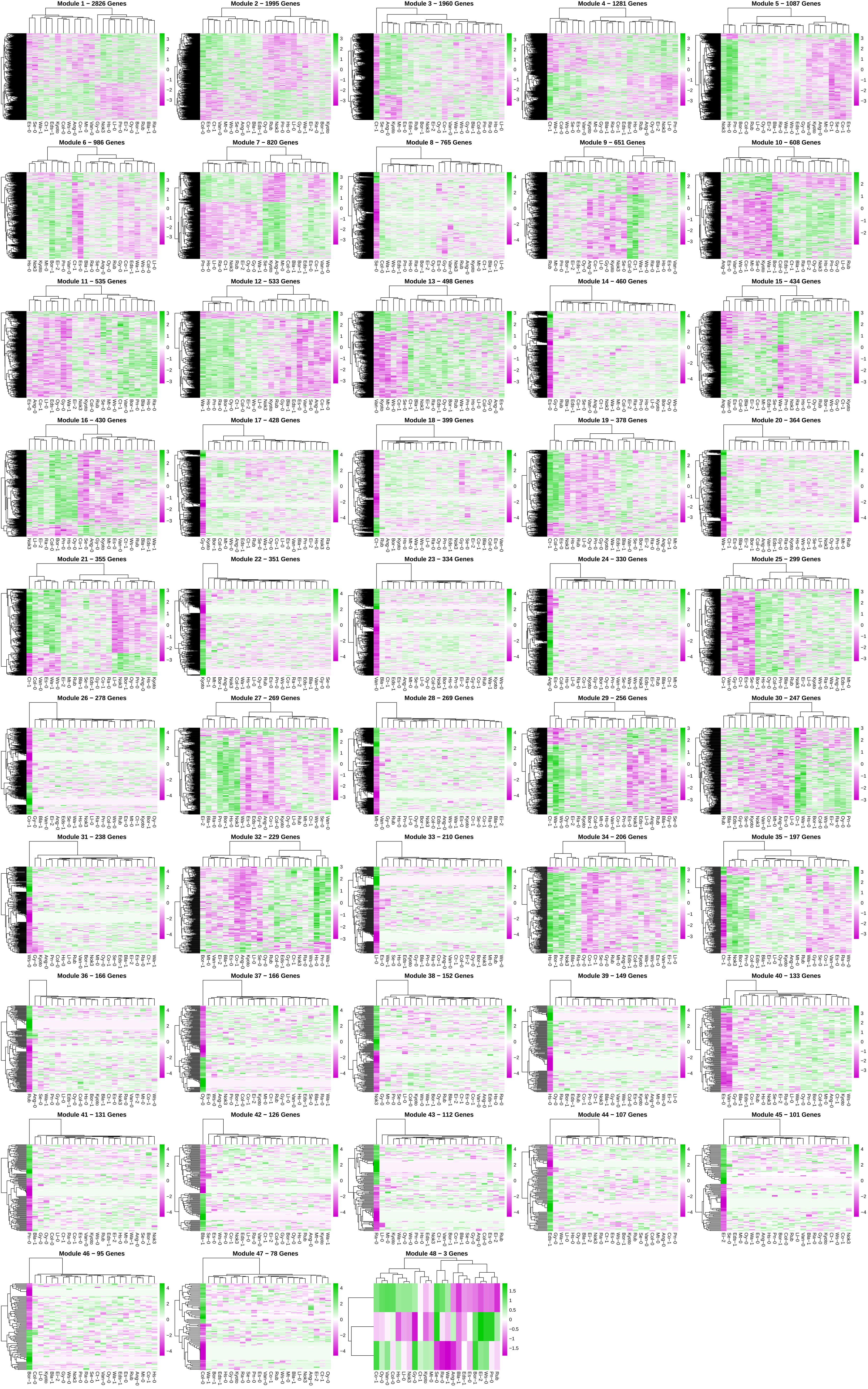
Module 1 – 2826 Genes
Module 2 – 1995 Genes
Module 3 – 1960 Genes
Module 4 – 1281 Genes
Module 5 – 1087 Genes
Module 6 – 986 Genes
Module 7 – 820 Genes
Module 8 – 765 Genes
Module 9 – 651 Genes
Module 10 – 608 Genes
Module 11 – 535 Genes
Module 12 – 533 Genes
Module 13 – 498 Genes
Module 14 – 460 Genes
Module 15 – 434 Genes
Module 16 – 430 Genes
Module 17 – 428 Genes
Module 18 – 399 Genes
Module 19 – 378 Genes
Module 20 – 364 Genes
Module 21 – 355 Genes
Module 22 – 351 Genes
Module 23 – 334 Genes
Module 24 – 330 Genes
Module 25 – 299 Genes
Module 26 – 278 Genes
Module 27 – 269 Genes
Module 28 – 269 Genes
Module 29 – 256 Genes
Module 30 – 247 Genes
Module 31 – 238 Genes
Module 32 – 229 Genes
Module 33 – 210 Genes
Module 34 – 206 Genes
Module 35 – 197 Genes
Module 36 – 166 Genes
Module 37 – 166 Genes
Module 38 – 152 Genes
Module 39 – 149 Genes
Module 40 – 133 Genes
Module 41 – 131 Genes
Module 42 – 126 Genes
Module 43 – 112 Genes
Module 44 – 107 Genes
Module 45 – 101 Genes
Module 46 – 95 Genes
Module 47 – 78 Genes
Module 48 – 3 Genes

**Supplemental Figure S6: Genes responding to *S. sclerotiorum* can be classified into at least 48 coexpression modules**.

Heatmap showing the expression profile of genes associated with co-expression modules identified by WGCNA across the 23 accessions. Green indicates LFC > 0, and purple indicates LFC < 0. The number of genes per module is labeled in the heatmap title.

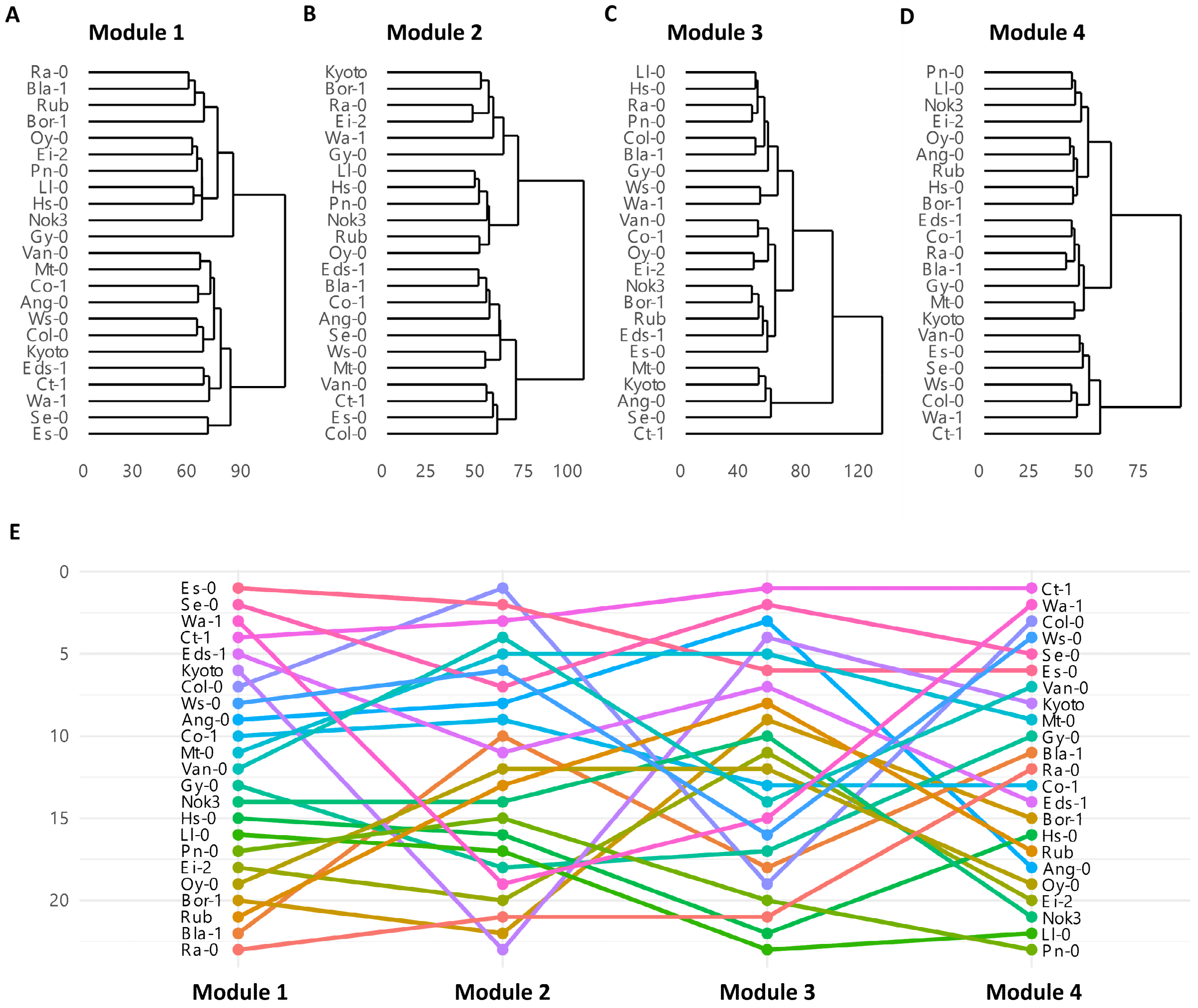


**Supplemental Figure S7: Transcriptome similarity among accessions varies across gene co-expression modules.**

**A, B, C, and D**. Dendrograms of accessions based on gene co-expression modules described in Figure S6. **E**. Comparison of the accession order derived from the dendrograms for each gene co-expression module.

.

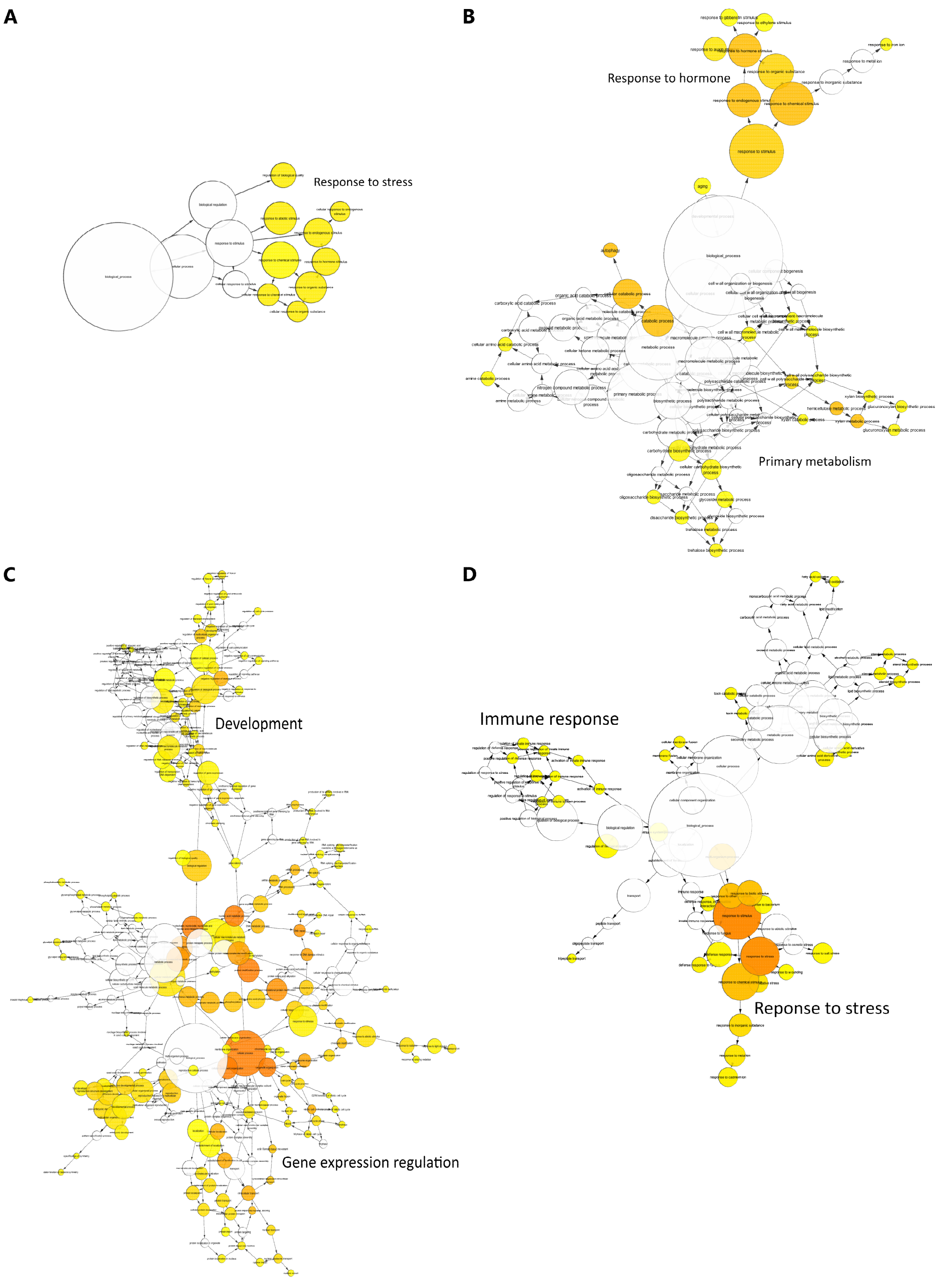


**Supplemental Figure S8: Gene ontology processes related to stress response, developmental processes, primary metabolism, and immune responses are enriched in co-expression modules 1 to 4.**

**A, B, C and D**. Gene ontology was obtained for the four larger modules (1 to 4) using the BINGO plugin from Cytoscape.

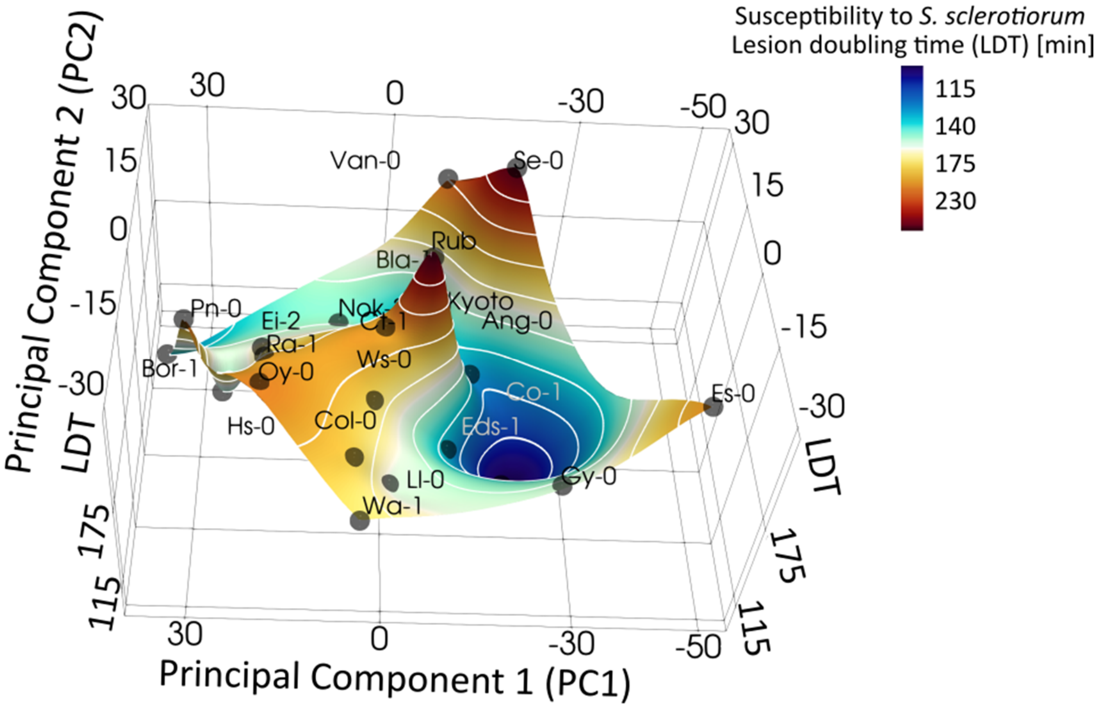


**Supplemental Figure S9: Transcriptome-resistance map showing accession proximity in the transcriptome space using only core differentially expressed genes.**

Transcriptome-resistance map showing accession proximity in the transcriptome space in response to *S. sclerotiorum* inoculation (X-Y plane) and resistance to *S. sclerotiorum* (Z-axis, color scale). The x and y axes represent components 1 and 2 of a principal component analysis using LFC data for core differentially expressed genes only. The color gradient represents disease resistance from more susceptible (blue) to more resistant (red), with experimentally studied accessions shown as small labelled spheres. Lines of equal resistance (white) are provided for legibility.
